# Rv2061c, a Putative F_420_-Dependent Oxidoreductase, is Required for Tolerance to Malachite Green and Other Antimicrobials and Oxidative Stress Protection in *Mycobacterium tuberculosis*

**DOI:** 10.64898/2026.10.08.757578

**Authors:** Dalin Rifat, Thomas R. Ioerger, Eric L. Nuermberger

## Abstract

The pathogenicity of *Mycobacterium tuberculosis* relies on its ability to withstand hostile host environments. Coenzyme F_420_ is critical for redox metabolism in *Mycobacterium tuberculosis*, an organism distinguished by its abundant use of this low-redox-potential cofactor and a repertoire of F_420_-dependent enzymes. While F_420_-dependent mechanisms are known to confer protection against antimicrobials and oxidative stress, the specific enzymes responsible remain largely unidentified. In this study, using the F_420_-dependent detoxification of malachite green as a phenotypic marker, we performed a genome-wide screen in *Mycobacterium tuberculosis* using transposon mutagenesis and whole genome sequencing. The results of transposon mutagenesis and whole genome sequencing revealed that, among the 28 genes encoding F_420_-dependent enzymes, only mutations in *fgd* (essential for F_420_H_2_ re-generation) and *Rv2061c*, a gene encoding a putative F_420_-dependent oxidoreductase, were highly susceptible to malachite green. Subsequent experiments confirmed that Rv2061c and its homolog MT2120 are required for detoxifying malachite green and crystal violet. Furthermore, the *Rv2061c* mutant exhibited increased susceptibility to isoniazid and pretomanid; as well as menadione-induced oxidative stress. We also identified mutations in the transcriptional repressor Rv0678 as a key determinant of resistance to malachite green and crystal violet. This study establishes Rv2061c as a pivotal F_420_-dependent oxidoreductase that protects *Mycobacterium tuberculosis* from malachite green and other antimicrobial compounds as well as oxidative stress and further supports Rv0678 as a master regulator of multidrug resistance. These findings expand our knowledge of *Mycobacterium tuberculosis* defense mechanisms and highlight F_420_-dependent pathways as a potential target for new drug development.

## Introduction

The deazaflavin cofactor F_420_[7,8-didemethyl(14–16)-8-hydroxy-5-deazariboflavin-5′-phosphoryllactyl(glutamyl)*_n_* glutamate] is abundant in mycobacteria and is also produced by archaea and other actinobacteria (1–9). Despite its structural similarity to flavin mononucleotide and flavin adenine dinucleotide (FAD), F_420_ is functionally analogous to the nicotinamide adenine dinucleotide cofactors NAD and NADP. As an obligate two-electron carrier with a low standard redox potential (-340 mV, compared to -219 mV for FAD and - 320 mV for NAD), F_420_ enables thermodynamically challenging reductions of otherwise recalcitrant organic molecules (10–13)(14–16). Although F_420_ is best known for its central role in methanogenesis, where it serves as a redox cofactor for CO_2_ reduction to methane in methanogenic *Euryarchaeota* (17), its importance in bacteria, though secondary to that of NAD and NADP, extends to numerous metabolic pathways (18, 19). These include the biodegradation of aflatoxins and nitrophenol, the biosynthesis of tetracycline and lincosamide antibiotics, and the reduction of biliverdin, quinones, and mycolic acids (10, 11, 14, 15, 20–22).

Despite its abundance and unique redox properties, the scope of F_420_^’^ s role in mycobacterial physiology is incompletely understood. Mycobacteria utilize F_420_-dependent redox mechanisms to survive harsh conditions, including oxidative and nitrosative stress as well as exposure to antimicrobial compounds (11, 19, 20, 23–26). For example, mutations in *fbiC* and *fbiD* genes, which are responsible for biosynthesis of F_420_ precursors F0 and F_420_-0, impair the growth and survival of *Mycobacterium tuberculosis* (Mtb) under oxidative and nitrosative stress as well as progressive hypoxia (11, 24). Studies in Mtb and *Mycobacterium smegmatis* have demonstrated F_420_H_2_-dependent tolerance to selected antimicrobials. For example, mutants defective in F_420_ biosynthesis or F_420_H_2_ generation are more susceptible to multiple TB drugs including isoniazid (INH), clofazimine (CFZ), moxifloxacin, linezolid (LZD), bedaquiline (BDQ), rifampin (RIF), ethambutol (EMB) and pyrazinamide (PZA), although the magnitude of the effect may vary between drugs and between mycobacterial species (11, 19, 24), and several recent studies have demonstrated that F_420_H_2_-deficient Mtb mutants may in fact be less susceptible to low inhibitory concentrations of CFZ (27–29). F_420_H_2_-deficient mutants are also more susceptible to malachite green (MG) (24, 25, 30), a triphenylmethane dye with antimicrobial properties (31) commonly used in mycobacterial media to prevent contamination by faster-growing microorganisms. At concentrations employed in Middlebrook 7H10 agar (0.25 mg/L), Middlebrook 7H11 agar (1 mg/L), and Löwenstein–Jensen (LJ) medium (400 mg/L), MG inhibits most bacteria and fungi but is tolerated by mycobacteria due to their F_420_-dependent detoxification capacity (13, 24, 25), converting MG to its non-toxic, colorless reduced form, leucomalachite green (11, 19, 24).

Mycobacterial genomes encode the largest putative protein family of F_420_-dependent enzymes (1–3, 14). For example, the genome of Mtb is predicted to encode at least 28 F_420_-dependent proteins, most of which remain functionally uncharacterized (32). While some have been shown to participate in stress responses and cell wall biosynthesis (11, 20, 23), the specific F_420_-dependent enzymes responsible for detoxifying antimicrobial compounds, including TB drugs and MG, are largely unknown. Their characterization could provide critical insights into TB drug detoxification and reveal novel therapeutic targets.

Comparative genomic analyses have identified three major families of F_420_-dependent enzymes in Mtb and other actinobacteria: luciferase-like monooxygenases (LLM), pyridoxamine 5′-phosphate oxidases (PPOX), and deazaflavin-dependent nitroreductases (DDN), indicating that F_420_-dependent proteins constitute the largest family in mycobacteria (14, 32). Of at least 28 distinct F_420_-dependent enzymes encoded by Mtb, only a few have been characterized (3, 11, 20, 33). One key enzyme is the F_420_-dependent glucose-6-phosphate dehydrogenase (Fgd, Rv0407) (3, 34–36). Mtb appears to depend solely on Fgd to catalyze the reduction of F_420_ to F_420_H_2,_ using glucose 6-phosphate, whose intracellular level is 17- to 130-fold higher in Mtb than in other bacteria (36). Disruption of *fgd* in *M. smegmatis* results in hypersusceptibility to nitrosative/oxidative stress and increased susceptibility to antimicrobials, including MG, RIF, EMB, INH, CFZ and PZA (19, 23). The F_420_H_2_-dependent nitroreductase Ddn (Rv3547) reductively activates the nitroimidazole prodrugs pretomanid and delamanid; and mutations in *fgd*, *ddn* or genes required for F_420_ biosynthesis confer resistance to these drugs (24, 32, 33). Although purified Ddn and its homologs Rv1261 and Rv1558 exhibit antioxidant activity *in vitro* and are proposed to function as F_420_H_2_-dependent quinone reductases (Fqrs) (11), their physiological roles require confirmation through gene deletion studies in intact Mtb. F_420_-dependent enzymes can be quite promiscuous in their activity. For example, six purified F_420_-dependent reductases from *M. smegmatis* were shown to reduce 10 of 20 tested antimicrobial compounds, including MG, RIF, and CFZ, but not INH (19). Despite this *in vitro* activity, the physiological role of these enzymes remains unconfirmed, and the functional equivalents in Mtb have yet to be identified.

Therefore, we employed MG detoxification as a phenotypic marker, in combination with transposon mutagenesis and whole genome sequencing (Tn-seq), to conduct a comprehensive genome-wide screen for F_420_-dependent enzymes responsible for antimicrobial detoxification in Mtb. In this study, we performed Tn-seq analysis on a Mtb H37Rv transposon mutant library exposed to MG, aiming to identify genes involved in the detoxification of MG, selected TB drugs, and oxidative stress. Among the 28 predicted F_420_-dependent proteins, Rv2061c, a putative F_420_-dependent oxidoreductase, emerged as a promising candidate. Functional characterization confirmed that Rv2061c is involved not only in the detoxification of MG and crystal violet (CV), another triphenylmethane dye, but also in conferring protection against the TB drugs INH and pretomanid, and the pro-oxidant compound menadione. We also confirmed that loss-of-function mutations in *Rv0678*, a known determinant of BDQ and CFZ resistance, also mediate resistance to MG.

## Results

### Tn-seq identifies F_420_-dependent genes required for Mtb survival on exposure to MG

We prepared 9 DNA libraries from purified genomic DNA of Mtb Tn mutant pools and performed deep sequencing. The deep sequencing data were used to calculate log_2_FC for each gene by comparing Tn junction read abundances between MG-exposed and control samples, followed by identification of conditionally-essential genes via resampling-based statistical analysis. Raw insertion counts at TA sites in each sample, mapped to the H37Rv (NC_000962.3) genome using TPP in Transit were shown in Dataset_1. Tn-seq results are presented in Dataset_2. We identified 44 and 24 genes for which Tn insertions were statistically significantly less abundant in mutant pools exposed to 3 and 30 µg/ml of MG, respectively, indicating loss of fitness (i.e., attenuated growth) with loss of gene function. Conversely, Tn insertions were significantly more abundant in 21 genes and 1 gene, respectively, indicating greater fitness (i.e., resistance) under these same conditions. Table 1 lists genes demonstrating the largest changes (log_2_FC > 3 or < - 3) following exposure to 3 or 30 µg/ml of MG. Two genes, *fgd* (*Rv0407*) and *caeA/hip1* (*Rv2224c*), showed consistent attenuation under both conditions. This result for *fgd* was consistent with previous findings that F_420_-dependent glucose-6-phosphate dehydrogenase (Fgd), the sole enzyme responsible for reducing F_420_ using glucose-6-phosphate in Mtb, is required for MG detoxification in mycobacteria (3, 34–36). Our analysis also identified insertions in *fbiC* (Rv1173), which is essential for F_420_ biosynthesis (25), as attenuating (log_2_FC= - 2.31) at 30 µg/ml of MG (Dataset_2), supporting the established role of F_420_ in MG detoxification. We hypothesized that one or more F_420_-dependent enzymes is responsible for detoxifying MG in Mtb. Among the 28 genes predicted to encode F_420_-dependent enzymes, only *fgd* and *Rv2061c* were markedly underrepresented among genes with Tn insertions at both MG concentrations (log_2_FC < -3), although *Rv2061c* did not reach statistical significance (adjusted *P* value > 0.05) (Table 2). This suggested *Rv2061c*, which encodes an uncharacterized F_420_-dependent oxidoreductase of the PPOX family, participates in MG detoxification.

**Table 1.** The list of significant genes with log_2_ FC >3 or <-3 after exposing to MG of 3 and 30 µg/ml.

| Gene ID | Name | Description | Sites | Log<sub>2</sub>FC after MG exposure |  |
| --- | --- | --- | --- | --- | --- |
| | | | | 3 μg/ml | 30 μg/ml |
| *Rv0472c* | *madR* | probable transcriptional regulatory protein (possibly TetR-family) | 10 | -4.05 |  |
| *Rv1836c* | - | hypothetical protein Rv1836c | 43 | -4.03 |  |
| *Rv3267* | *lcp1* | conserved hypothetical protein (CpsA-related protein) | 23 | -3.98 |  |
| *Rv2190c* | *ripC* | hypothetical protein Rv2190c | 20 | -3.89 |  |
| *Rv2224c* | *hip1, caeA* | probable exported protease serine protease Hip1(hydrolase important for pathogenesis) | 29 | -3.43 | -2.09 |
| *Rv0007* | - | possible conserved membrane protein | 10 | -3.35 |  |
| *Rv1420* | *uvrC* | excinuclease ABC subunit C | 29 | -3.33 |  |
| *Rv2696c* | - | conserved hypothetical alanine and glycine and valine rich protein | 10 | -3.33 |  |
| *Rv0929* | *pstC2* | phosphate-transport integral membrane ABC transporter PstC2 | 17 | -3.29 |  |
| *Rv0407* | *fgd* | probable F<sub>420</sub>-dependent glucose-6-phosphate dehydrogenase Fgd | 20 | -3.21 | -3.78 |
| *Rv1821* | *secA2* | preprotein translocase subunit SecA | 39 | -3.07 |  |
| *Rv0928* | *pstS3* | periplasmic phosphate-binding lipoprotein<br>PstS3 (Pbp-3) (PstS3) (PhoS1) | 22 | -3.06 |  |
| *Rv0999* | - | hypothetical protein Rv0999 | 11 | -3.01 |  |
| *Rv2198c* | *mmpS3* | probable conserved membrane protein<br>MmpS3 | 14 |  | -3.93 |
| *Rv0998* | - | lysine acetyltransferase (Mt-Pat) | 13 |  | -3.77 |
| *Rv2051c* | *ppm1* | polyprenol-monophosphomannose synthase<br>Ppm1 | 33 |  | -3.41 |
| *Rv2091c* | - | probable membrane protein | 19 |  | -3.37 |
| *Rv1823* | - | hypothetical protein Rv1823 | 15 |  | -3.29 |
| *Rv3421c* | - | hypothetical protein Rv3421c | 12 | 3.44 |  |
| *Rv3422c* | - | hypothetical protein Rv3422c | 6 | 4.55 |  |
| *Rv3804c* | *fbpA* | secreted antigen 85A FbpA (mycolyl<br>transferase 85A)<br>(fibronectin-binding protein A) (antigen 85<br>complex A) | 16 |  | 3.88 |

**Table 2.**
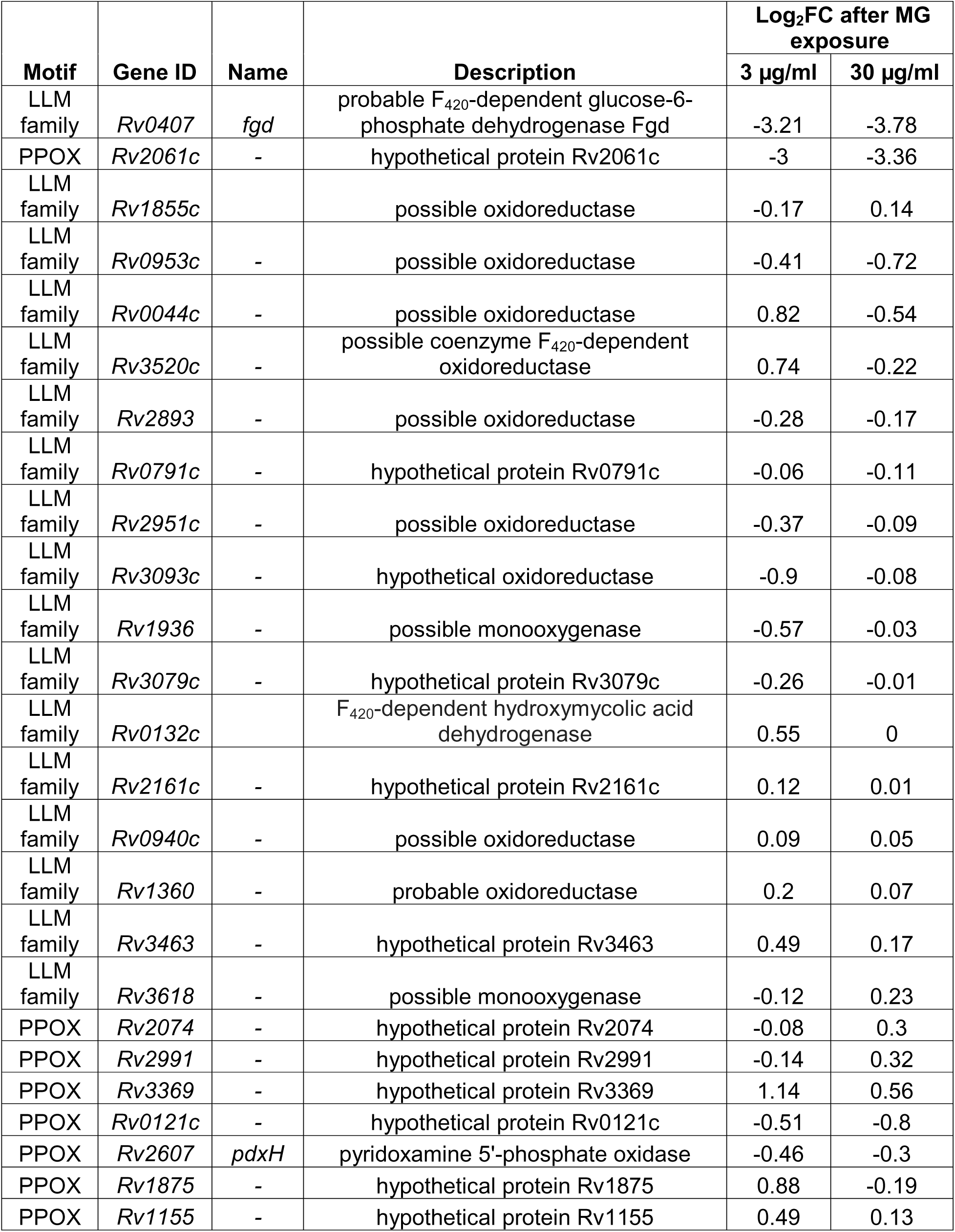
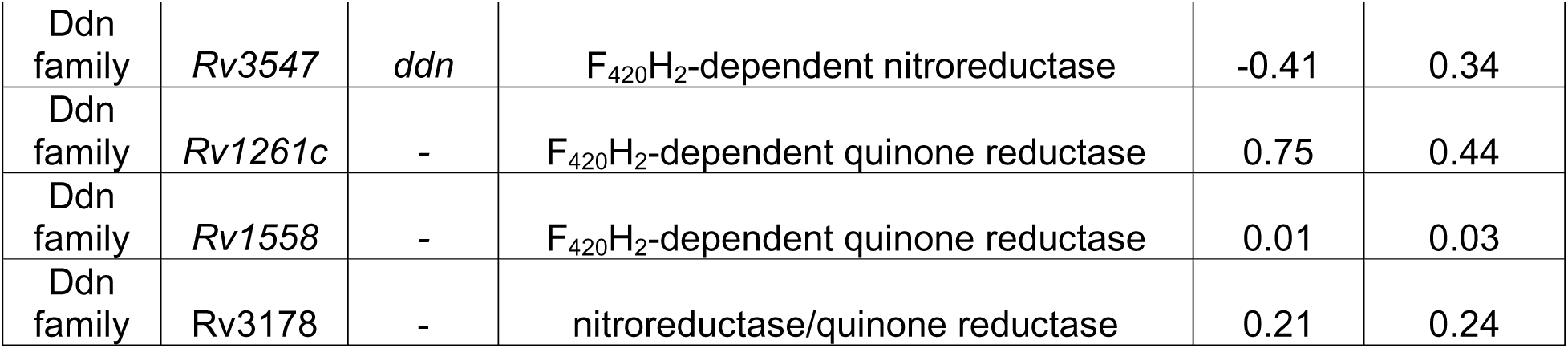
Tn-seq results of predicted genes that encode F_420_-dependent proteins in Mtb after exposure to MG.

### Rv2061c (or MT2120) is required for detoxification of MG and CV in Mtb

To confirm that *Rv2061c* is required for MG detoxification, we retrieved a mutant, POA_B20, previously selected from a wild-type Mtb H37Rv background, that contains a premature stop codon mutation at C39G (Tyr13*) in *Rv2061c* (38). This mutant was complemented with the wild-type *Rv2061c* gene after determining from our Tn-seq results that its flanking genes appeared irrelevant to MG susceptibility (Dataset_2). Notably, although the open reading frames of *Rv2061c* and *MT2120* (ortholog in Mtb strain CDC1551) are annotated as 405 and 444-bp, respectively, DNA sequence alignment revealed that the 444-bp of *MT 2120* region and its intergenic sequences flanking the two adjacent genes are identical to the 405-bp of *Rv2061c* region and its corresponding intergenic sequences with its neighboring genes. To verify the role of MT2120 in MG susceptibility, we retrieved another mutant, HG1504, with a Tn insertion in *MT2120* and complemented it with the wild-type *MT2120* allele. Since the POA_B20 mutant also carries a mutation at A1106G (His369Arg) in *clpC1* conferring pyrazinoic acid resistance (38), we included an additional mutant, POA_B21, which was selected contemporaneously with POA_B20 but contains only a *clpC1* mutation, to better control for any effect of *clpC1* mutations on MG detoxification. We then determined the MICs of MG and CV for the mutant, complemented, and wild-type strains. As shown in Table 3, mutations in *MT2120* and *Rv2061c* were associated with lower MG MICs of 10 and 20 µg/ml, compared to MG MICs of 20 and 80 µg/ml for their respective wild-type strains. Similarly, the MIC of CV was 4 times lower against the *MT2120* mutant compared to the wild-type MIC. Complementation with the respective wild-type genes restored the wild-type MIC levels or, presumably due to overexpression from the *hsp60* promoter, resulted in even higher MICs for both compounds (Table 3). This phenotypic susceptibility was corroborated by a second assay demonstrating the decolorization of MG and CV in cultures of wild-type and complemented strains exposed to sub-MIC concentrations, but not in cultures of the *Rv2061c* and *MT2120* mutants exposed to the same concentrations of the compounds (Figure 1). The hypersusceptibility phenotypes of the mutants were further confirmed by spot assays on 7H9 agar plates containing increasing concentrations of MG and CV. After 28 days of incubation at 37°C, the *Rv2061c* and *MT2120* mutants showed impaired growth compared to the control strains as the MG or CV concentration increased (Figures 2A and 2B). Both MIC values of MG and CV against POA_B21 (isolated *clpC1* mutant) and its growth on MG- or CV-containing 7H9 agar plates suggested that the *clpC1* mutation was not responsible for the hypersusceptibility of POA_B20 to MG and CV, as its phenotype was similar to that of the wildtype H37Rv (Table 3 and Figure 2).

**Figure 1.**
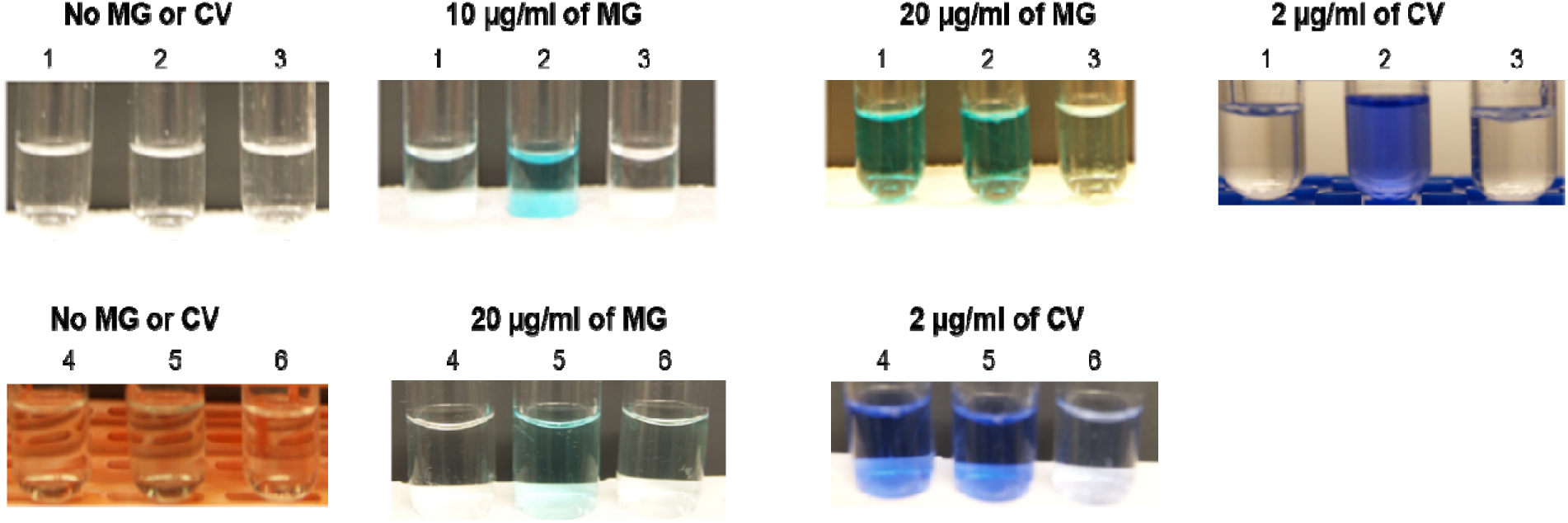
Rv2061c and MT2120 are involved in detoxification of MG and CV. Minimum inhibitory concentration (MIC) tests were performed against MG and CV using Mtb strains. Bacterial suspensions were prepared in 7H9 broth (without Tween-80) at approximately 10^5^ CFU/ml in 4 ml round-bottom tubes. The tubes were incubated at 37°C statically for 14 days. After the incubation period, photographs were taken of tubes containing critical concentration of the dyes: 10 and 20 µg/ml for MG and 2 µg/ml for CV. The strains tested were as follows: 1. CDC1551; 2. HG1504 (*MT2120* mutant); 3. Complemented strain of HG1504; 4. H37Rv; 5. POA_B20 (*Rv2061c* mutant); 6. Complemented strain of POA_B20.

**Figure 2.**
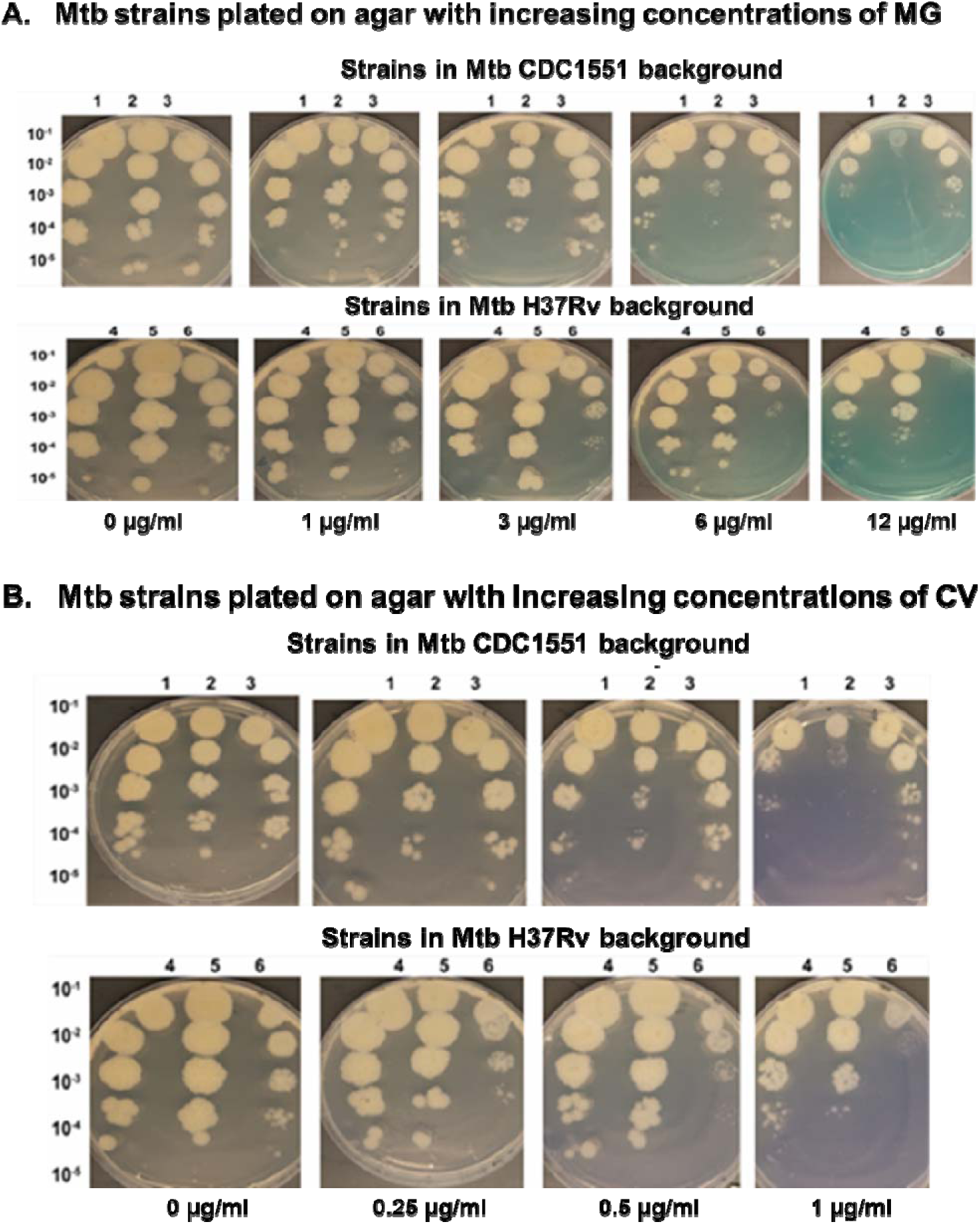
Impaired growth of the *Rv2061c* and *MT2120* mutants on 7H9 agar plates with increasing concentrations of MG and CV compared to the control strains. Following growth to mid-log phase, Mtb strains were subjected to serial 10-fold dilutions. Aliquots (10 µl) of each dilution were spotted in order of decreasing bacterial concentration, ranging from 10^-1^ to 10^-5^, onto 7H9-agar plates containing varying concentrations of MG (A) or CV (B). To enable direct comparison, strains from the same Mtb strain background were spotted on the same plates and incubated at 37°C for 28 days. Strains in Mtb CDC1551 background included: 1. CDC1551; 2. HG1504 (*MT2120* Tn mutant); 3. Complemented strain of HG1504. Strains in Mtb H37Rv background included: 4. H37Rv; 5. POA_B21 (*clpC1* mutant); 6. POA_B20 (*Rv2061c* mutant).

**Table 3.** MIC results of the *Rv2061c* (or *MT2120*) mutant and the *Rv0678* mutant alongside their respective wild-type and complemented strains against MG and CV.

| Strain | MIC (µg/ml) |  |
| --- | --- | --- |
|  | MG | CV |
| **H37Rv** | **40-80** | **2** |
| **POA_B20 (*Rv2061c*: C39G(Tyr13\*); *clpC1*: A1106G(His369Arg); *Rv2402*: C1135A (Arg379Ser))** | **20** | **2** |
| **complemented strain of POA_B20 (pMH94-*hsp60*promoter-*Rv2061c*)** | **80** | **4** |
| **POA_B21 (*clpC1*: G296A(Gly99Asp))** | **40** | **4** |
| **CDC1551** | **20** | **4** |
| **HG1504 (POI at *MT2120*: 319/444-bp)** | **10** | **1** |
| **complemented strain of HG1504 (pMH94-*hsp60*promoter-*MT2120*)** | **80** | **8** |
| ***Rv0678* mutant (ins. A at POI 275/498-bp)** | **>80** | **4** |
| **Complemented strain of *Rv0678* mutant (pMH94-*hsp60*promoter-*Rv0678*)** | **40** | **2** |

### Mutations in *Rv2061c* in Mtb increase susceptibility to isoniazid and pretomanid

To determine whether Rv2061c and MT2120 contribute to intrinsic resistance to TB drugs, we first performed broth microdilution MIC assays using mutant and control strains against a panel of drugs, including INH, BDQ, CFZ, LZD, RIF, and pretomanid. No significant differences greater than one dilution in MIC were observed between the mutants and their respective wild-type strains (data not shown). Given the limited sensitivity of MIC assays, we further investigated drug susceptibility using spot assays on 7H11 agar plates containing increasing concentrations of the drugs. While no growth differences were observed for the *MT2120* mutant compared to wild-type CDC1551 on plates containing INH (Figure 3), or on CFZ or LZD (data not shown), the *Rv2061c* mutant exhibited a clear growth defect on 0.06 µg/ml of INH (Figure 3). This increased susceptibility to INH was not attributable to a secondary *clpC1* mutation, as the POA_B21 control strain with an isolated *clpC1* mutation displayed a wild-type phenotype (Figure 3). However, complementation of the *Rv2061c* mutant failed to fully restore the wild-type growth phenotype on agar containing INH (data not shown). To quantitatively validate these findings, we performed time-kill assays in 7H9 broth. Under INH pressure (0.06 µg/ml), the *Rv2061c* mutant showed a significant reduction in viability compared to wild-type H37Rv, with log_10_CFU/ml reductions of -3.5 versus -2.8 at day 4 (*p*=0.01) and -5.5 versus -2.4 at day 8 (*p*=0.04) (Figure 4A). Complementation with *Rv2061c* successfully restored wild-type levels of INH tolerance (*p*=0.006). We also assessed the role of Rv2061c in susceptibility to pretomanid, a prodrug requiring activation by the F_420_-dependent nitroreductase Ddn for TB activity (33). The *Rv2061c* mutant was significantly more susceptible to pretomanid than wild-type H37Rv at day 4 (*p*=0.006); and this phenotype was successfully complemented (*p*=0.006) (Figure 4B). While the mutant and the wild type converged by a later time point, the complemented strain overexpressing *Rv2061c* was clearly less susceptible to pretomanid, avoiding the CFU decline seen in the other strains (Figure 4B).

**Figure 3.**
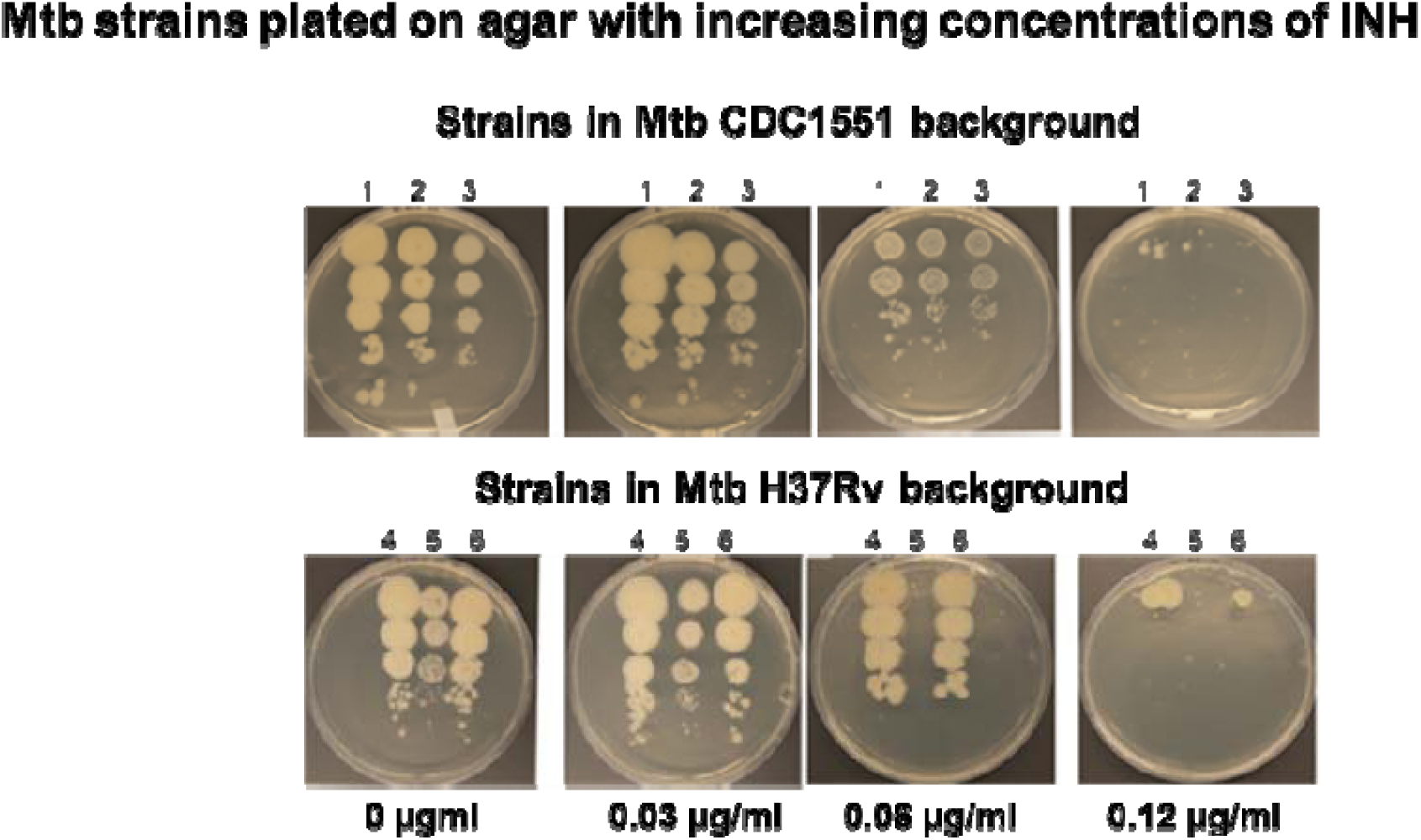
A significant growth defect on INH-containing plates was observed in a Mtb strain with a mutation in *Rv2061c*, not in one with a mutation in *MT2120*. Following growth to mid-log phase, Mtb strains were subjected to serial 10-fold dilutions. Aliquots (10 µl) of each dilution were spotted in order of decreasing bacterial concentration, ranging from 10^-1^ to 10 ^-5^ (from top to bottom), onto 7H11 agar plates containing varying concentrations of INH. To enable direct comparison, a group of strains was spotted on the same plates and incubated at 37°C for 28 days. The Mtb strains included in the study were as follows: 1. CDC1551; 2. HG1504 (*MT2120* Tn mutant); 3. Complemented strain of HG1504; 4. H37Rv; 5. POA_B20 (*Rv2061c* mutant); 6. POA_B21 (*clpC1* mutant).

**Figure 4.**
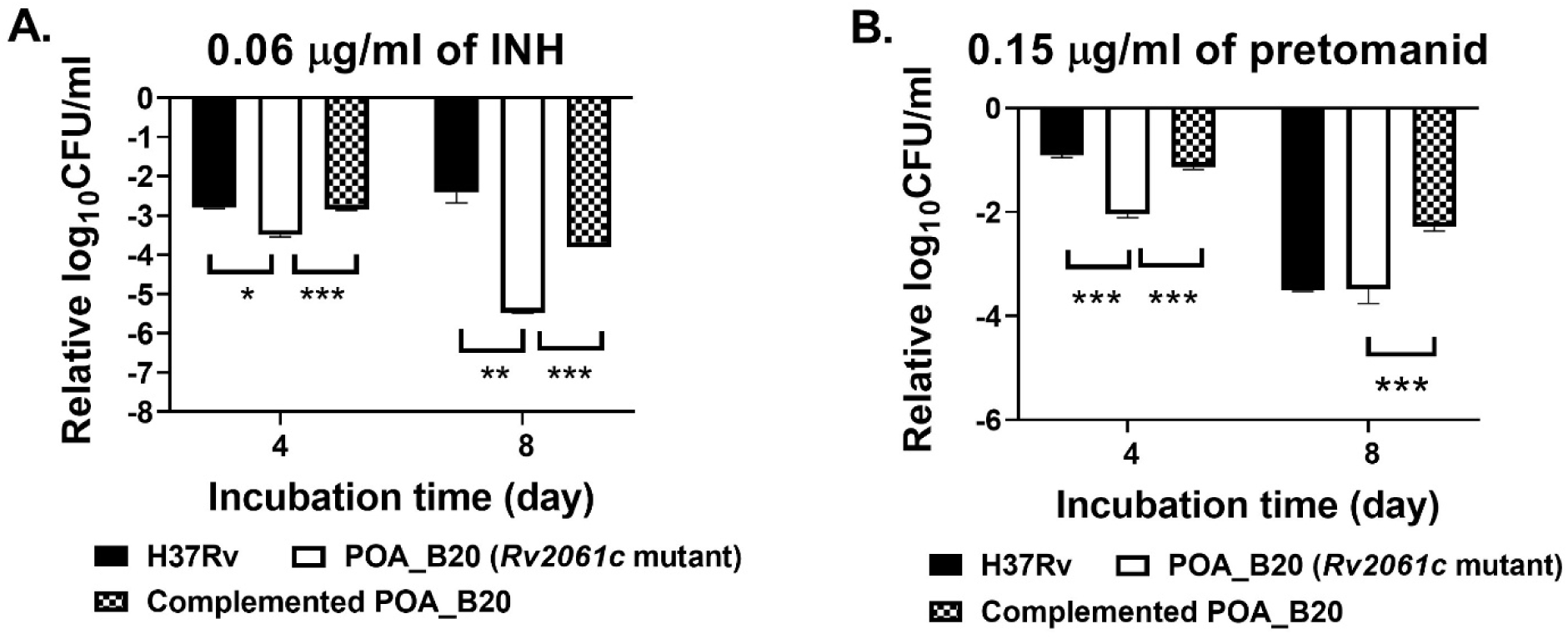
Mutations in *Rv2061c* resulted in greater susceptibility to INH and pretomanid. Mtb strains were cultured in 7H9 broth containing different concentrations of drugs for 8 days, including no drug controls. Viability was assessed by plating aliquots on Day 0 (before the treatment started) and after 4, and 8 days of drug exposure, respectively. The relative log_10_ CFU/ml (drug-treated vs. untreated control) was calculated for each strain at Day 4 and Day 8, respectively. A. Mtb strains exposed to 0.06 µg/ml of INH; B. Mtb strains exposed to 0.15 µg/ml of pretomanid. \**p* = 0.01; ** *p* = 0.04; *** *p* = 0.006.

### The survival of Mtb under oxidative stress was impaired by the absence of Rv2061c

To determine if Rv2061c and MT2120 are involved in survival under oxidative stress, H37Rv and CDC1551 wild-type strains, along with their respective mutants carrying mutations in *Rv2061c*, *MT2120*, *ddn*, and *clpC1*, as well as their complemented strains, were exposed to various concentrations of menadione (0, 20, 50, and 100 µM) for up to 10 days. The change in log_10_ CFU/ml (menadione-treated vs. untreated culture) for each strain was calculated at day 3, day 7, and day 10. Although no or slight reductions in CFU were observed for all H37Rv-derived strains after 3 days of exposure to 20 µM menadione, the *Rv2061c* mutant (POA_B20) experienced a >3 log_10_ CFU/ml reduction in CFU/ml compared to untreated samples at day 7 (*p*=0.002 vs. H37Rv), which was partially restored by complementation (*p*=0.016 vs. H37Rv) (Figure 5A). Similar results were observed after 10 days of incubation under 20 µM menadione indicating that *Rv2061c* is required for survival under oxidative stress. The impaired survival under oxidative stress was not attributable to the *clpC1* mutation, as the *clpC1* mutant (POA_B21) exhibited survival comparable to that of H37Rv upon menadione exposure. Nonetheless, the roles of the and *Rv2402* mutations in survival of the POA_B20 mutant under oxidative stress may require further investigation. The *ddn* mutant (KA091) behaved similarly to the wild-type, suggesting that, unlike Rv2061c, Ddn (F_420_-dependent nitroreductase) is not essential under oxidative stress (Figure 5A). H37Rv and H37Rv-derived mutants carrying mutations in *Rv2061c*, *ddn*, and *clpC1* experienced nearly complete loss of viability at higher menadione concentrations, including 100 µM menadione at day 3 and 50 µM menadione at day 7 (data not shown). Interestingly, CDC1551-derived mutants adapted to 20 µM menadione as well as the wild-type CDC1551 strain across all time points (Figure 5B); and also was not more susceptible than the wild-type CDC1551 parent to 50 and 100 µM menadione, indicating that neither MT2120 nor MT3561 (Ddn) is required for survival under these oxidative stress conditions (data not shown).

**Figure 5.**
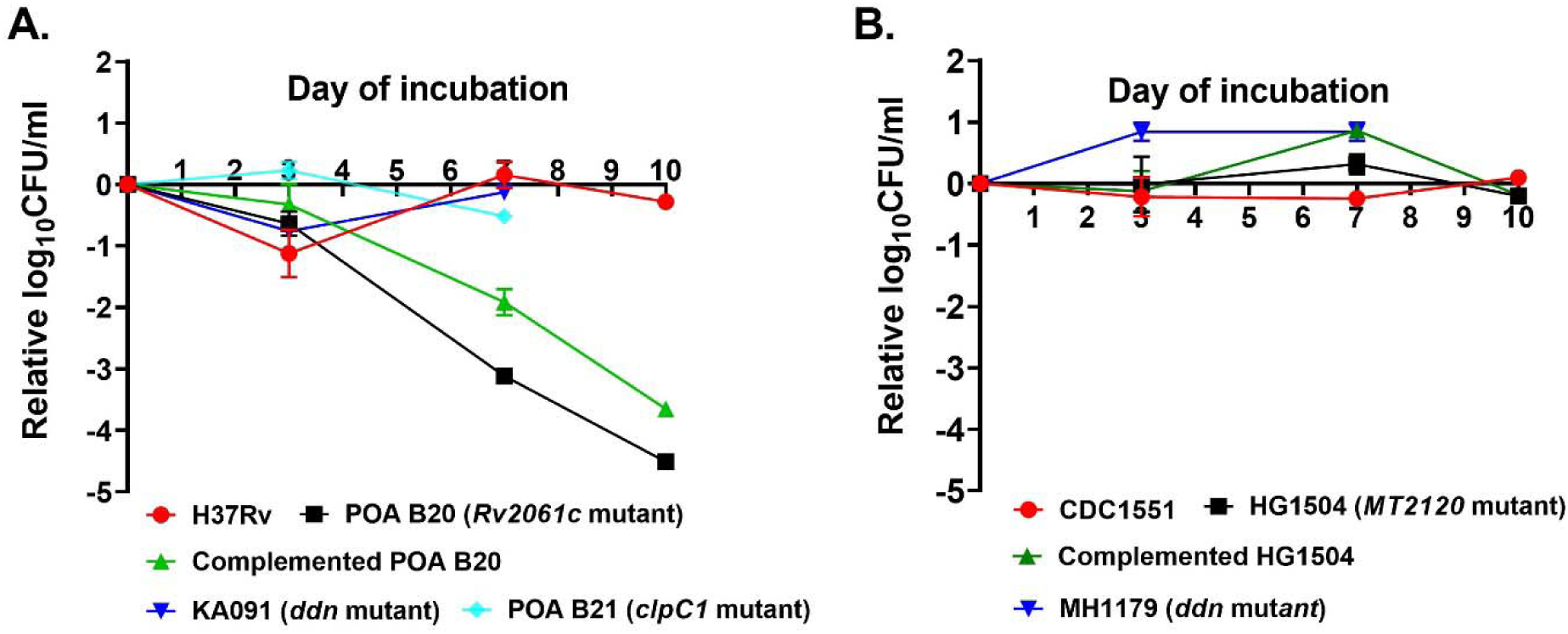
Survival of Mtb under menadione-induced oxidative stress was impaired by the absence of Rv2061c in the H37Rv strain background. Mtb strains were cultured in 7H9 broth containing 0 and 20 µM menadione for 10 days. Viability was assessed by plating aliquots on Day 0 (immediately after treatment initiation), Day 3, Day 7, and Day 10. The relative log_10_ CFU/ml (menadione-treated vs. untreated control) was calculated for each strain at Day 3, Day 7, and Day 10, respectively. A. Relative log_10_ CFU/ml of Mtb H37Rv-derived strains; B. Relative log_10_ CFU/ml of Mtb CDC1551-derived strains.

### Mutations in *Rv0678* associated with bedaquiline and clofazimine resistance also confer resistance to MG and CV

Our Tn-seq analysis also identified genes in addition to *Rv2061c* and F_420_-related genes in which Tn insertions attenuated survival on MG, including genes encoding transcriptional regulators, transporters, and cell wall synthesis components (Dataset_2). Notably, Tn insertions in *mmpL5* (*Rv0676c*) attenuated survival at both MG concentrations (adjusted *P* < 0.05). In Mtb, *mmpL5* and *mmpS5* (*Rv0677c*) form an operon encoding an efflux transporter regulated by the downstream transcriptional regulator MmpR5 (Rv0678) (46, 47). Loss-of-function mutations in *Rv0678* confer resistance to BDQ and CFZ by derepressing *mmpL5* and *mmpS5* expression (48–50). Our Tn-seq analysis of Mtb exposed to 30 µg/ml of MG revealed a log_2_FC of 1.78 for *Rv0678*, though this did not reach statistical significance (Dataset_2). To validate genes naturally associated with MG resistance, we cultured the Mtb H37Rv Tn mutant pool on 7H9 agar plates containing 60 and 120 µg/ml of MG. A colony selected on 120 µg/ml of MG was found to carry a Tn insertion in *Rv0678* at position 435/498-bp, consistent with our Tn-seq finding. The resistance phenotype of this mutant was further confirmed by plating it along with the following strains at similar bacterial density on 7H9 agar plates containing MG at concentrations ranging from 0 to 120 µg/ml: wild-type H37Rv; a previously described isogenic BDQ-resistant strain with an insertion mutation in *Rv0678* (A inserted at 275/498-bp) (39); B101 (an *fbiD* [*Rv2983*] mutant) (24); and two CDC1551-derived *mmpL5* Tn-insertion mutants (STN0592 and STN1424). After 28 days of incubation all strains grew similarly on plates without MG (data not shown). In contrast to the wild-type H37Rv strain, the *Rv0678* mutant exhibited resistance to MG at concentrations up to 120 µg/ml (Figure 6). The MG-hypersusceptible *fbiD* (B101) mutant (24), along with the two *mmpL5* Tn mutants, showed a growth defect relative to wild-type H37Rv (Figure 6), which is consistent with our Tn-seq results as described above, given that both H37Rv and CDC1551 showed similar growth on MG-containing plates (Figure 2). These two *mmpL5* mutants are also hypersusceptible to BDQ relative to their parent strain CDC1551 (data not shown). To further validate our findings, we complemented the isogenic BDQ-resistant *Rv0678* mutant strain with wild-type *Rv0678*. The MIC results confirmed that mutations in *Rv0678* conferred resistance to MG and CV; and introduction of wild-type *Rv0678* into the mutant fully restored the wild-type phenotype (Table 3).

**Figure 6.**
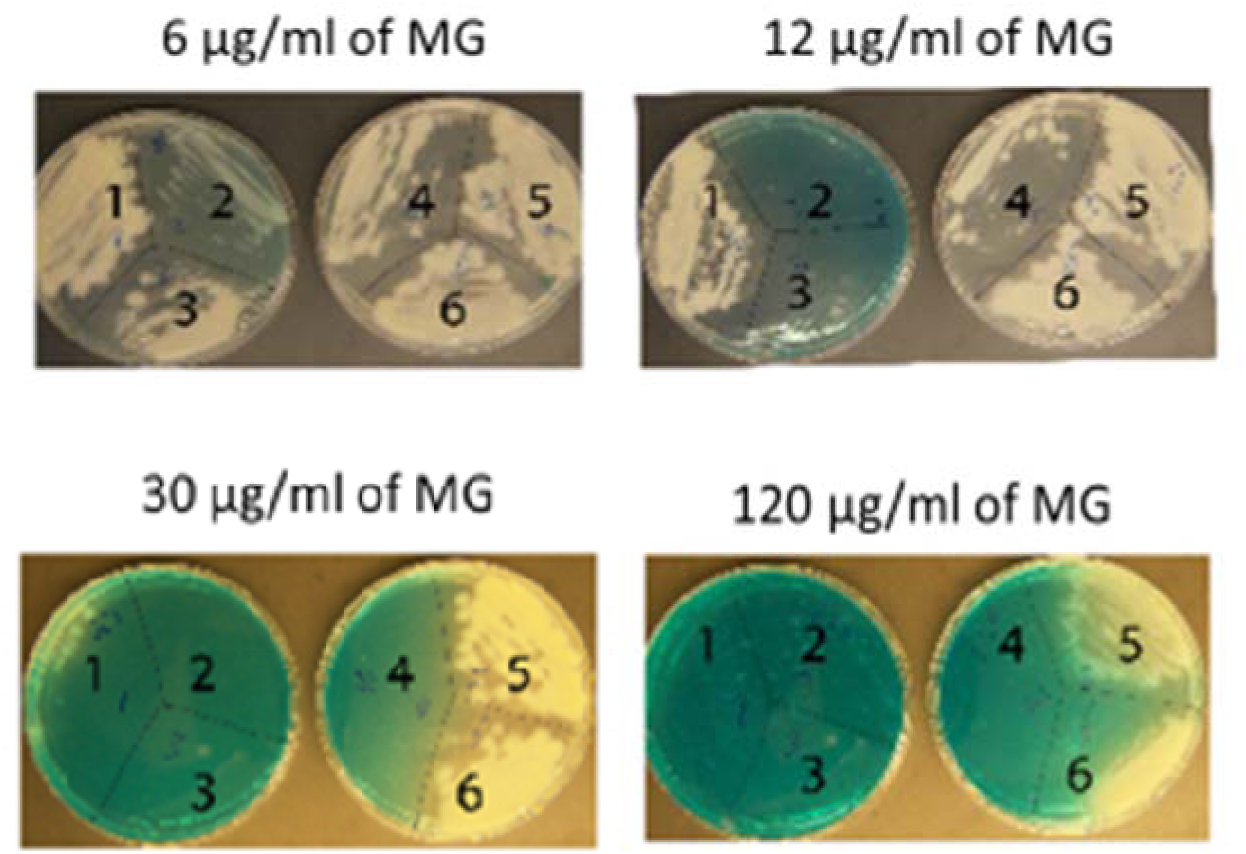
Mutations In *Rv0678* and *MT0705* (*Rv0676c*) conferred resistance and hypersusceptibility to MG, respectively. Mtb strains were cultured in 7H9 broth to mid-log phase and diluted in PBS to same O.D. _600nm_. Bacterial suspensions were streaked onto 7H9-agar plates containing various concentrations of MG and incubated at 37°C for 28 days. The strains used were as follows: 1. H37Rv; 2. B101 (*Rv2983* mutant); 3. STN0592 (*MT0705* mutant); 4. STN1424 (*MT0705* mutant); 5. Isogenic *Rv0678* mutant (ins. A at 274_275-bp); 6. *Rv0678* Tn mutant (POI:435/498-bp, isolated from this study).

## Discussion

The ability of Mtb to persist within the host and survive antibiotic therapy poses a major challenge to effective treatment of TB. This resilience of Mtb is attributable in part to its diverse mechanisms for withstanding harsh environments. The low-redox-potential coenzyme F_420_ and its dependent pathways have garnered significant attention for their unique roles in mycobacterial redox metabolism under stress. In this study, we conducted a genome-wide Tn-seq screen of Mtb exposed to MG and identified Rv2061c, a putative F_420_-dependent oxidoreductase, as an enzyme required for Mtb tolerance to MG and CV, the TB drugs INH and pretomanid, as well as menadione-induced oxidative stress.

Mycobacteria have abundant F_420_ and F_420_-dependent enzymes. For example, Mtb and *M. smegmatis* are predicted to have 28 and 76 F_420_-dependent enzymes, respectively, suggesting a unique reliance on an F_420_-assisted redox lifestyle (9, 14, 32). Despite F_420_‘s proven roles in protection against stress conditions and antimicrobials, the specific F_420_-dependent enzymes responsible remain largely unidentified. We therefore used susceptibility to MG, an antimicrobial compound detoxified by mycobacteria in an F_420_H_2_-dependent manner, as a phenotypic marker to screen for these enzymes in Mtb. Our Tn-seq analysis revealed that, among the 28 genes predicted to encode F_420_-dependent enzymes in Mtb, only Tn insertions in *fgd* and *Rv2061c* led to significant attenuation under MG exposure. As *fgd* encodes the F_420_-dependent glucose-6-phosphate dehydrogenase essential for reducing F_420_ to F_420_H_2_, this finding underscores its critical role in F_420_-dependent detoxification systems (35). Follow-up studies in two Mtb strains (H37Rv and CDC1551) confirmed that loss of Rv2061c (or its ortholog MT2120), a PPOX class F_420_-dependent oxidoreductase, impaired MG decolorization and growth in the presence of MG, suggesting it is the dominant F_420_-dependent enzyme responsible for MG detoxification in Mtb. Although other F_420_-dependent enzymes may contribute to detoxification of MG, they could not be identified using our methodology. Importantly, loss-of-function mutations in the F_420_-dependent nitroreductase Ddn, which confer resistance to pretomanid and delamanid, did not confer hypersusceptibility to MG in our Tn-seq screen, which further supports our previous finding that defects in F_420_ biosynthesis and reduction to F_420_H_2_ reduce the ability to recover Mtb on MG-containing media, while mutations in *ddn* do not (24).

MG is a synthetic dye used in textiles, leather and paper goods, and also used illegally in many parts of the world as an antimicrobial/antiparasitic in aquaculture because of its low cost, ready availability, and high efficacy against pathogens of fish and other aquatic species (51). However, MG is toxic to mammalian cells and has therefore been banned by the U.S. Food and Drug Administration (52). The degradation of this recalcitrant dye has been studied in many microorganisms, including mycobacteria, primarily for the treatment of dye-contaminated wastewater and to mitigate environmental and public health risks (13, 53–56). To date, only a limited number of enzymes have been reported to degrade MG through oxidative mechanisms, including lignin peroxidase (LiP) and laccase from white-rot fungi and birds’ nest fungi (56) and manganese peroxidase (MnP) from *Pseudomonas aeruginosa*, and *Enterobacter roggenkampii* (53), respectively; or through NADH-dependent reductive mechanisms, such as triphenylmethane reductase from *Citrobacter* sp. (55). Mycobacteria are intrinsically resistant to MG (13), suggesting that enzymes from mycobacteria may be promising candidates for MG biodegradation. An early study of MG decolorization in *Mycobacterium avium* strain A5 showed that the membrane fraction exhibited at least a 5-fold higher specific decolorization rate than did the crude extract, where the reduced form of MG, leucomalachite green, was detected; this activity was significantly inhibited by boiling or treatment with 2,4-dinitrophenol, an inhibitor of proton translocation, and was not enhanced by the addition of NADH or NADPH (13). Consistent with this observation, Rv2061c has been confirmed to be a membrane-associated protein and is predicted to require the coenzyme factor F_420_H_2_ (32, 57, 58). Others reported that metyrapone, an inhibitor of cytochrome P450 enzymes, suppressed MG decolorization (13), suggesting a possible role for cytochrome P450 enzymes in this process (13). However, this hypothesis is not supported by our Tn-seq results, as no significant changes in log_2_ FC values were observed for the 20 genes encoding cytochrome P450 proteins in Mtb (ranging from - 0.64 to 0.66; Dataset_2) (59). However, the possibility that multiple such enzymes contribute to a redundant capacity for detoxification cannot be excluded by our results. Homology analysis of the Rv2061c protein sequence across mycobacterial species using NCBI reveals a high degree of conservation with some mycobacteria. For example, Rv2061c shares approximately 80% identity with members of *M. avium* complex, 69% identity with MSMEG_3863 in *M. smegmatis* MC^2^155, and 51% identity with MAB_2203 in *Mycobacterium abscessus subsp. abscessus* ATCC 19977. The MG degradation pathway is complex and involves multiple reaction steps, as proposed for *Exiguobacterium sp.* MG2 (54). Whether cytochrome P450 enzymes or other enzymes participate in MG decolorization in Mtb remains to be determined.

Beyond F_420_-dependent enzymes, our Tn-seq results also revealed other genes of interest associated with tolerance to the effect of MG. For example, *Rv2224c*, encoding the cell envelope-associated serine protease known as CaeA (carboxylesterase A) or Hip1 (hydrolase important for pathogenesis)(60, 61), was required for growth on media containing ≥3 µg/ml of MG. Loss-of-function mutations in *Rv2224c* enhance host innate immune responses, impair Mtb survival in macrophages, and increase susceptibility to multiple TB drugs, including INH, RIF, carbapenems, ethambutol, and clarithromycin, supporting its interest as a potential drug target (60–64). We also noted substantial overlap between other genes with Tn insertions sensitizing Mtb to MG in our study (Table 1) and those found to sensitize Mtb to vancomycin, RIF, INH, EMB and/or meropenem in a previous Tn-seq study (e.g., *mmpS3*, *secA2*, *lcp1*, *ppm1*, *Rv1836c*, *madR* (*Rv0472c*), *Rv0007*) (63). Like Xu et al, we hypothesize that the functional common denominator among these mutants is likely to be increased permeability of the Mtb cell envelope to antimicrobials and other toxic molecules. We note that Xu et al used 7H9 media (which does not contain MG) for antibiotic selection of Tn libraries. However, because MG is ubiquitous in solid media that may be used to cultivate Mtb for Tn-seq and other similar screening methodologies, selection of Tn mutants or conditional knockdown strains with chemical stresses in the presence of MG could bias results towards the genes associated with differential susceptibility to MG that we have identified here. In this context, our gene list in Table 1 may serve as an important reference set for potential false positive hits related to differential MG susceptibility.

Additionally, our study has newly identified de-repression of the *mmpL5-mmpS5* operon via loss-of-function mutations in *Rv0678* as a novel determinant of MG resistance. Mutations in *Rv0678* (*mmpR5*) have garnered intense interest owing to their established role in BDQ and CFZ resistance (46, 47), as well as resistance to novel DprE1 inhibitors in clinical development (65–67). Indeed, CFZ exposure was found to up-regulate expression of *Rv2061c* over 3 times (log2 FC=1.749) in DNA microarray studies (68). Thus, our results provide further evidence that such mutations confer low-level resistance to a broad spectrum of antimicrobials, presumably through enhanced efflux via the *mmpS5-mmpL5* transporter.

The pathogenicity of Mtb relies on its ability to combat hostile host environments. Following inhalation and phagocytosis by alveolar macrophages, Mtb encounters oxidative, nitrosative, and acidic stress, as well as nutrient limitation (69–72), and later faces hypoxia within necrotic granulomas (73). In addition to known pathways for detoxifying free radicals, such as those involving catalase, superoxide dismutase, DosR and WhiB3 regulons (74–78), F_420_-dependent redox mechanisms also aid survival under these conditions (9, 11, 23, 32). We identified Rv2061c as the first F_420_-dependent enzyme, aside from Fgd, shown to protect Mtb from both antimicrobial compounds and oxidative stress. Our finding that the *Rv2061c* mutant was attenuated in the presence of menadione supports a model in which Rv2061c is able to catalyze F_420_H_2_-dependent two-electron reduction of quinones to quinols, thereby competing with NADH-dependent quinone reductases to prevent one-electron reduction to cytotoxic semiquinone radicals and superoxide through redox cycling in the presence of oxygen because of the low redox potential of F_420_(11). Mutants of *MT2120* as well as *Rv2061c* (during early exposure) showed better survival than wild-type strains, possibly indicating pre-adaptation due to the mutations. However, the *Rv2061c* mutant became hypersusceptible with prolonged incubation, confirming its role in longer-term adaptation to oxidative stress and suggesting that a longer exposure might be needed to reveal the role of MT2120. Notably, we also demonstrated that Ddn is not required for menadione resistance in Mtb, despite a prior study demonstrating F_420_-dependent quinone reductase activity by purified Ddn and its homologs Rv1261 and Rv1558 (11). This discrepancy may arise from the redundant functions of these enzymes or other differences between activity in a purified system and essentiality for physiological function in whole cells. For example, many F_420_-dependent oxidoreductases may exhibit F_420_-dependent quinone reductase activity when tested in isolation, but Rv2061c may assume the dominant share of this activity in whole cells. The relevance of Rv2061c is further underscored by its significant upregulation in granulomas and peri-cavitary lung tissue from human TB samples compared to *in vitro* cultures (76). While our data establish its role in oxidative stress, further study is needed to determine if Rv2061c is required under other conditions such as nitrosative stress and hypoxia.

Studies have established that the coenzyme F_420_ is essential for mycobacterial detoxification of furanocoumarins (e.g., methoxsalen), arylmethanes (e.g., MG and CV), and quinone analogues (e.g., menadione) (11, 24). While F_420_’s role in detoxifying MG, CV, and menadione is clear, its loss confers only a slight increase in susceptibility to selected clinical TB drugs in *M. smegmatis* based on MIC data (32). Extending this investigation to Rv2061c, we found that although the mutant showed no significant MIC differences, it exhibited significant growth and survival defects in CFU-based assays when exposed to INH and pretomanid. The apparent relatively broad substrate specificity of Rv2061c aligns with the known promiscuity of F_420_-dependent enzymes. For instance, six purified F_420_H_2_-dependent reductases from *M. smegmatis* catalyze the reduction of diverse compounds, including menadione, MG, RIF, and CFZ (19). Notably, none of those enzymes reduced INH. This may be due to the fact that the *M. smegmatis* ortholog of Rv2061c, MSMEG_3863, was not characterized in the prior study (19)(28). The susceptibility of the *Rv2061c* mutant could be attributed to either the loss of direct reductase activity against these drugs or the loss of a general antioxidant property. The latter is particularly relevant for INH, a prodrug activated by KatG, whose enzymatic activity is induced under oxidative stress (74). Supporting this, a comprehensive genetic screen in a murine TB model found that *Rv2061c* Tn insertion mutants were significantly depleted (log_2_FC = - 4.26, *p*= 0.0007) after INH treatment (79). Interestingly, while the prodrug pretomanid requires F_420_-dependent activation by Ddn for its anti-TB activity (33), we found that the detoxifying enzyme Rv2061c contributed to early-stage protection, a role that was eventually overcome by the activating pathway.

Finally, we observed strain-dependent variations in drug and menadione response between H37Rv and CDC1551 in the absence of Rv2061c and MT2120. For example, the mutation in *Rv2061c*, but not a mutation in MT2120, increased sensitivity to INH and reduced survival under menadione-induced oxidative stress. Given that the annotated open reading frame and the flanking regions of *Rv2061c* are identical to those of *MT2120* in terms of DNA sequence, we speculate that the phenotypic variation stems from heterogeneity in the genetic background of their parent strains, such as strain-specific regulatory mechanisms and protein interaction networks, rather than from differences in the locus itself. Although a complementation strategy was used, potential effects from additional mutations in the *Rv2061c* mutant (POA_B20) cannot be ruled out. Therefore, generating a clean *Rv2061c* deletion mutant is warranted to verify the phenotypic differences between Rv2061c and MT2120 and to more fully characterize their detailed responses to a broad spectrum of TB drugs and other stresses. Rv2061c is annotated as a member of the PPOX family, a prediction supported by SIMBAL (sites inferred by metabolic background assertion labeling) analysis, which yielded a high score indicative of an F_420_-binding protein (32). However, its F_420_-dependent reductase activity has not yet been biochemically verified *in vitro*. Our findings on Rv2061c help to address a critical knowledge gap in mycobacterial stress metabolism and highlight its potential as a target for TB drug development. In addition, the potential application of purified Rv2061c for the biodegradation of environmental MG warrants further investigation.

## Materials and Methods

### Bacteria, media, antimicrobial compounds, and reagents

The Mtb H37Rv Tn mutant library was generously provided by Dr. Petros Karakousis (37). Wildtype strains H37Rv and CDC1551 were obtained from our laboratory stock stored at - 80°C (24). The following isogenic mutant strains were derived from the parental H37Rv strain: POA_B20 (*Rv2061c*: C39G(Tyr13*); *Rv3596c* (*clpC1*): A1106G(His369Arg); *Rv2402*: C1135A (Arg379Ser)); POA_B21 (*clpC1*: G296A(Gly99Asp)) (38); a *Rv0678* (*mmpR5*) mutant (A inserted at POI 275/498-bp) (39); KA091 (*Rv3547* (*ddn*): IS6110 ins. in D108) (24); and B101 (*Rv2983* (*fbiD*): Ala198Pro) (24). Additionally, several Himar1 Tn insertion mutants were derived from the parental CDC1551 strain with POI (position of Tn insertion) as follows: HG1504 (POI at *MT2120*: 319/444-bp); MH1179 (POI at *MT3651* (*ddn*): 52/456-bp); STN1424 (POI at MT0705 (*mmpL5*): 1540/2895); and STN0592 (POI at MT0705 (*mmpL5*): 830/2895) (40). For bacterial cultivation, we used Difco Middlebrook 7H9 broth (BD) supplemented with 10% oleic acid-albumin-dextrose-catalase (OADC) complex (BD), 0.2% glycerol, and 0.05% Tween 80 (Sigma-Aldrich), unless otherwise specified. Antimicrobial susceptibility testing employed 7H9 broth without Tween 80. Tn-seq studies utilized 7H9 agar plates containing Middlebrook 7H9 broth (BD), 0.2% glycerol, and 1.5% Bacto agar (BD) (24). Routine cultivation and drug testing were performed on BBL 7H11 agar base (BD) supplemented with 10% OADC and 0.5% glycerol. MG oxalate was obtained from Alfa Aesar. Pretomanid, BDQ, and LZD were provided by the TB Alliance (New York, NY). INH, CFZ, RIF, CV, and menadione were purchased from Sigma-Aldrich. All compounds were freshly prepared in dimethyl sulfoxide except INH was dissolved in water.

### MG exposure of the Mtb Tn mutant library and whole genome sequencing analysis

The Mtb Tn mutant library was subcultured in 7H9 broth with shaking at 37°C until reaching an O.D._600nm_ of 1.0. The culture was then plated in triplicate on 7H9 agar plates containing either 3 µg/ml of MG, 30 µg/ml of MG, or no MG (control). Following 28 days of incubation at 37°C, mutant lawns were collected from the plates for genomic DNA purification using PureLink^TM^ Microbiome DNA purification Kit (Invitrogen). DNA libraries were prepared from purified genomic DNAs as previously described (41, 42), with primers listed in Table S1. Libraries were sequenced on an Illumina HiSeq platform generating 4.9 (3.5-9.6) million 150-bp paired-end reads per sample. Raw reads were processed using TPP in TRANSIT (43) and mapped to TA dinucleotide sites in Mtb H37Rv reference genome sequence (accession number NC_000962.3). The raw insertion counts were normalized using the Beta-Geometric Correction (BGC) in Transit (44), to mitigate the impact of read-count distributions that were highly skewed for some samples. For each gene with a Tn insertion, a log_2_ fold change (FC) in insertion abundance was calculated for MG-containing plates (3 and 30 µg/ml) relative to the MG-free control. A negative FC value indicates increased susceptibility of Mtb to MG when insertions occur in the gene, whereas a positive FC value indicates reduced susceptibility to MG when insertions occur. Resampling statistical analysis was used to determine when insertions were significantly more or less abundant in the MG-exposed Tn mutant libraries relative to the MG-unexposed libraries, using adjusted *P* values < 0.05 to define statistical significance (42).

### Selection of Tn mutant strains resistant to MG

The Mtb Tn mutant library was subcultured in 7H9 broth with shaking at 37°C until reaching an O.D._600nm_ of 1.0. The culture was then plated on 7H9 agar plates containing either 60 or 120 µg/ml of MG. After 28 days of incubation at 37°C colonies were selected from these plates and subcultured for DNA purification. Genomic DNA of individual colonies was used to determine the location of Tn insertions on the genome as previously described (40), with some modifications. Briefly, DNA was first digested with AluI (NEB, New England Biolabs) at 37°C for 1 hour followed by DNA cleaning, and then ligated to an annealed adaptor (AdTop/AdBtm) using Blunt/TA ligase Mater Mix (NEB) at 25°C for 30 min. To amplify the junction of Tn insertion regions, a Tn primer T7-A and an adaptor primer AP1 was used in PCR reactions with Q5 hot start high fidelity 2X master mix (NEB). The PCR products were purified prior to DNA sequencing. The sequences of the adaptors and the primers are listed in Table S1.

### Complementation of *Rv2061c*, *MT2120* and *Rv0678* in their corresponding mutants

To generate complemented strains, Mtb mutants carrying mutations in *Rv2061c*, *Rv0678*, and *MT2120* were transformed with *Mycobacterium-Escherichia coli* shuttle vector pMH94 (hygromycin resistant), which contained the Mtb *hsp60* promoter and the respective gene’s open reading frame (ORF). Briefly, the *hsp60* promoter and each ORF (*Rv2061c*, *Rv0678*, or *MT2120*) was PCR-amplified from Mtb H37Rv or CDC1551 genomic DNA using primers listed in Table S1. The PCR products were then ligated into XbaI-digested pMH94 using the NEBuilder HiFi DNA assembly kit (New England BioLabs) and transformed into *E. coli* competent cells. Plasmid constructs (pMH94-*hsp60*promoter-*Rv2061c*, pMH94-*hsp60*promoter-*Rv0678*, and pMH94-*hsp60*promoter-MT2120) were verified by DNA sequencing. These recombinant plasmids were electroporated into the respective mutant strains, and transformants were selected on 7H11 agar plates containing 50 μg/ml of hygromycin. Complemented strains were genetically confirmed by PCR-amplification of the hygromycin resistance gene and the insert junction regions from genomic DNA, using primers listed in Table S2. All of the mutant strains were successfully complemented (data not shown).

### MIC determination

A broth macrodilution assay was used to determine the MIC of MG and CV, while a broth microdilution assay was used for TB drugs (24, 45). For the macrodilution assay, mid-log-phase cultures of Mtb were adjusted to achieve a bacterial density of approximately 10^5^ colony-forming units (CFU)/ml in 7H9 broth without Tween 80 before 2.4 ml of the culture was added in 14 ml conical tubes, followed by addition of 100 µl of two-fold serially diluted solutions of MG or CV to reach final concentrations ranging from 5 to 320 µg/ml for MG and from 0.25 to 16 µg/ml for CV (24). The cultures were incubated at 37°C for 14 days without shaking, and the MIC was determined as the lowest concentration that prevented bacterial growth (i.e., no visible pellets observed at the bottom of the tubes). The color change of media containing MG or CV was documented on Day 14. For the microdilution assay, 96-well flat bottom sterile plates (Greiner) were used. First, 100 µl of 7H9 broth without Tween 80 was added to each well. Then, 100 µl of a drug solution diluted with 7H9 broth without Tween 80 was added to the first row of wells, followed by two-fold serial dilution across subsequent rows. Finally, 100 µl of the adjusted bacterial culture (prepared as described above) was added into each well. After 7 days of incubation at 37°C, 32.5 µl of alamarBlue (Thermo Fisher Scientific) was added to each well and plates were incubated 16-19 hours at 37°C in the dark. Fluorescence was measured using a BMG Optima microplate reader at 544nM excitation/590nM emission (45). Percentage inhibition was calculated as follows: the average fluorescence intensity of the no drug control wells was subtracted from the fluorescence intensity of the experimental wells, and their difference was divided by the average control and then multiplied by 100 times. The MIC was defined as the lowest drug concentration that resulted in at least 90% inhibition. Samples were prepared in triplicate and the experiment was repeated once under the same conditions.

### Antimicrobial growth inhibition assays

Mtb was grown in 7H9 broth to mid-log phase (O.D._600nm_ = 1.0). The cultures were then diluted to an O.D._600nm_ of 0.3 and subjected to 10-fold serial dilutions in sterile PBS, ranging from 10^-1^ to 10^-5^. 7H9 agar plates were prepared as previously described (24), containing varying concentrations of MG or CV. 7H11 agar was autoclaved, cooled to 50°C, and supplemented with antimicrobial compounds at specified concentrations before pouring into Petri dishes. Serial dilutions were spotted in 10 µl aliquots from top-to-bottom in columns in order of decreasing bacterial concentrations, with different strains arranged column-wise per plate. Plates were air-dried, then incubated at 37°C up to 28 days. Growth kinetics of Mtb strains were analyzed by documenting images of bacterial colonies on the plates.

### Time-kill assay in 7H9 broth

Mtb was cultivated in 7H9 broth to an O.D._600nm_ of 0.8 - 1. The cultures were adjusted to a bacterial density of approximately 10^6^ CFU/ml in 7H9 broth with Tween 80. TB drugs or menadione was added to these cultures to achieve the desired concentrations, respectively. Prior to addition, an aliquot of the cell suspension was serially diluted and plated on 7H11 agar for CFU counts. The drug-exposed cell suspensions, including no-drug controls, were then incubated for up to 10 days at 37°C with agitation. At different time points, aliquots of the cultures were removed, and serial 10-fold dilutions were plated on 7H11 agar. Plates were incubated for 28 days at 37°C before CFU counting. The change in log_10_CFU/ml was calculated by subtracting the CFU count of the no-drug controls from that of the drug-exposed samples at different time points, respectively, for each strain at each concentration of drug or menadione. Samples were prepared in duplicate, and student’s t-test was used to compare the mean change in CFU count between the strains tested.

## Supporting information

Dataset_1 Raw Tn-seq data

Dataset_2 Log2 fold change

## Supplemental materials

Dataset_1. Raw Tn-seq data from Mtb Tn mutant pools after exposure to varying concentrations of MG

Dataset_2. Log_2_FC in Mtb Tn mutant pools following exposure to MG relative to the no MG control

## Acknowledgement

We gratefully acknowledge funding support from the National Institutes of Health (UM1-AI179699 and R01-AI111992).

---

**Table S1.**
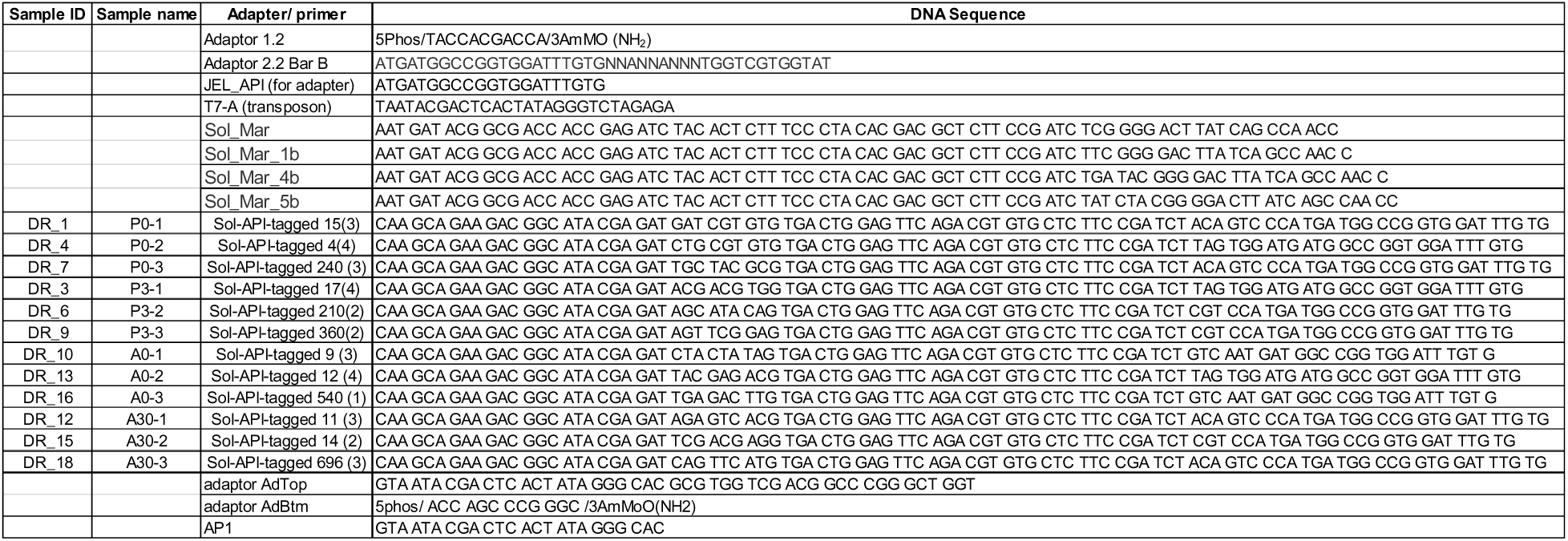
List of sequences of adapters and sequencing primers for DNA library preparation.

**Table S2.**
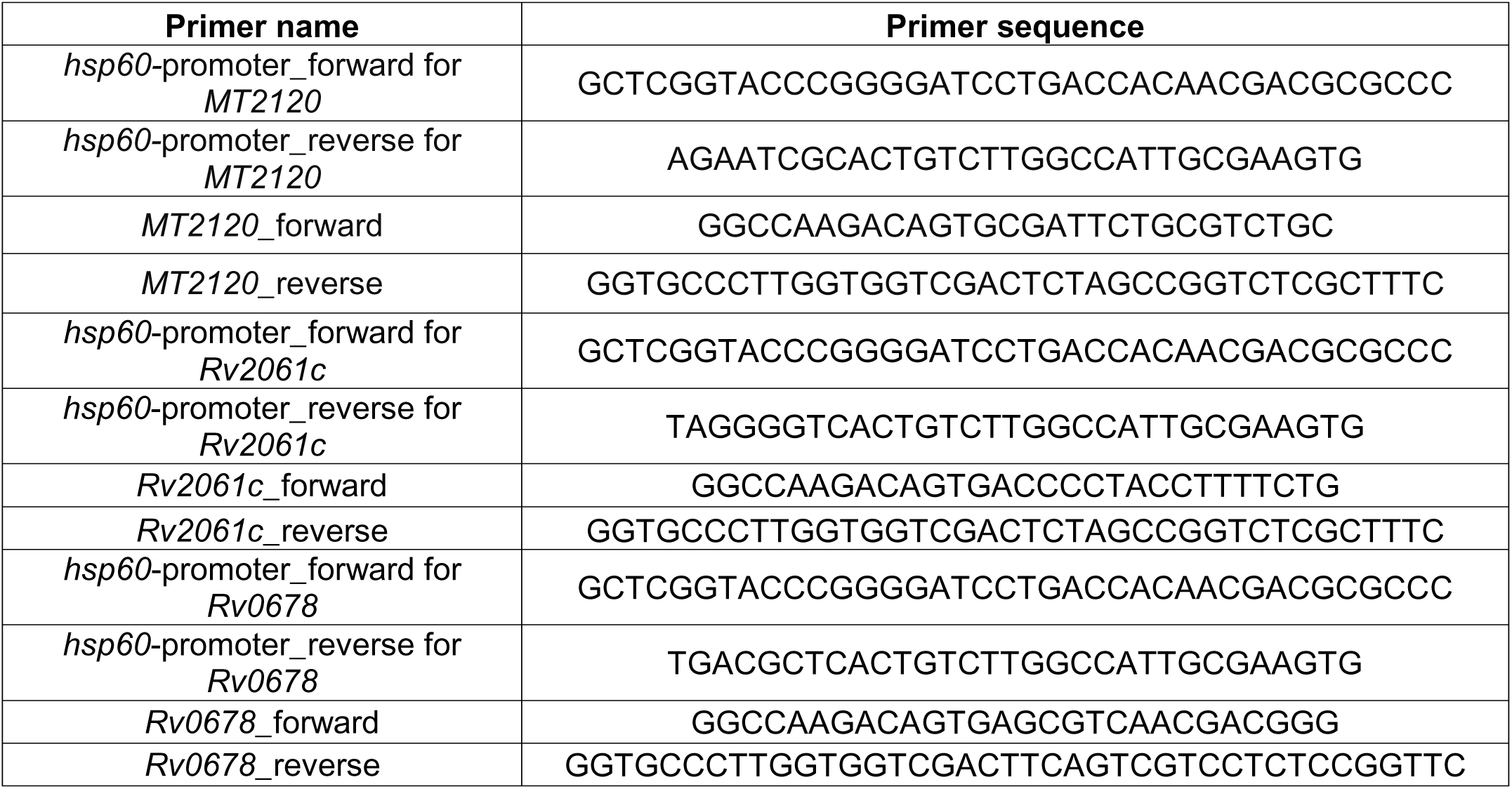
List of primers used for complementation of Mtb mutant strains.

## References

1. Naraoka T, Momoi K, Fukasawa K, Goto M. 1984. Isolation and identification of a naturally occurring 7, 8-didemethyl-8-hydroxy-5-deazariboflavin derivative from *Mycobacterium avium*. Biochimica et Biophysica Acta (BBA) - General Subjects 797:377–380.

2. Daniels L, Bakhiet N, Harmon K. 1985. Widespread Distribution of a 5-deazaflavin Cofactor in Actinomyces and Related Bacteria. Systematic and Applied Microbiology 6:12–17.

3. Purwantini E, Gillis TP, Daniels L. 1997. Presence of F420-dependent glucose-6-phosphate dehydrogenase in Mycobacterium and Nocardia species, but absence from Streptomyces and Corynebacterium species and methanogenic Archaea. FEMS Microbiol Lett 146:129– 134.

4. Cheeseman P, Toms-Wood A, Wolfe RS. 1972. Isolation and properties of a fluorescent compound, factor 420 , from Methanobacterium strain M.o.H. J Bacteriol 112:527–531.

5. Evans PN, Parks DH, Chadwick GL, Robbins SJ, Orphan VJ, Golding SD, Tyson GW. 2015. Methane metabolism in the archaeal phylum Bathyarchaeota revealed by genome-centric metagenomics. Science 350:434–438.

6. Kerou M, Offre P, Valledor L, Abby SS, Melcher M, Nagler M, Weckwerth W, Schleper C. 2016. Proteomics and comparative genomics of Nitrososphaera viennensis reveal the core genome and adaptations of archaeal ammonia oxidizers. Proc Natl Acad Sci U S A 113:E7937–E7946.

7. Ney B, Ahmed FH, Carere CR, Biswas A, Warden AC, Morales SE, Pandey G, Watt SJ, Oakeshott JG, Taylor MC, Stott MB, Jackson CJ, Greening C. 2017. The methanogenic redox cofactor F420 is widely synthesized by aerobic soil bacteria. ISME J 11:125–137.

8. Braga D, Last D, Hasan M, Guo H, Leichnitz D, Uzum Z, Richter I, Schalk F, Beemelmanns C, Hertweck C, Lackner G. 2019. Metabolic Pathway Rerouting in Paraburkholderia rhizoxinica Evolved Long-Overlooked Derivatives of Coenzyme F420. ACS Chem Biol 14:2088–2094.

9. Isabelle D, Simpson DR, Daniels L. 2002. Large-scale production of coenzyme F420-5,6 by using Mycobacterium smegmatis. Appl Environ Microbiol 68:5750–5755.

10. Taylor MC, Jackson CJ, Tattersall DB, French N, Peat TS, Newman J, Briggs LJ, Lapalikar GV, Campbell PM, Scott C, Russell RJ, Oakeshott JG. 2010. Identification and characterization of two families of F420H2-dependent reductases from Mycobacteria that catalyse aflatoxin degradation. Mol Microbiol 78:561–575.

11. Gurumurthy M, Rao M, Mukherjee T, Rao SPS, Boshoff HI, Dick T, Barry CE, Manjunatha UH. 2013. A novel F420-dependent anti-oxidant mechanism protects Mycobacterium tuberculosis against oxidative stress and bactericidal agents. Mol Microbiol 87:744–755.

12. Lapalikar GV, Taylor MC, Warden AC, Scott C, Russell RJ, Oakeshott JG. 2012. F420H2-Dependent Degradation of Aflatoxin and other Furanocoumarins Is Widespread throughout the Actinomycetales. PLOS ONE 7:e30114.

13. Jones JJ, Falkinham III JO. 2003. Decolorization of Malachite Green and Crystal Violet by Waterborne Pathogenic Mycobacteria. Antimicrob Agents Chemother 47:2323–2326.

14. Ahmed FH, Carr PD, Lee BM, Afriat-Jurnou L, Mohamed AE, Hong N-S, Flanagan J, Taylor MC, Greening C, Jackson CJ. 2015. Sequence–Structure–Function Classification of a Catalytically Diverse Oxidoreductase Superfamily in Mycobacteria. Journal of Molecular Biology 427:3554–3571.

15. Ebert S, Rieger PG, Knackmuss HJ. 1999. Function of coenzyme F420 in aerobic catabolism of 2,4, 6-trinitrophenol and 2,4-dinitrophenol by Nocardioides simplex FJ2-1A. J Bacteriol 181:2669–2674.

16. Li W, Khullar A, Chou S, Sacramo A, Gerratana B. 2009. Biosynthesis of Sibiromycin, a Potent Antitumor Antibiotic. Appl Environ Microbiol 75:2869–2878.

17. Thauer RK, Kaster A-K, Seedorf H, Buckel W, Hedderich R. 2008. Methanogenic archaea: ecologically relevant differences in energy conservation. Nat Rev Microbiol 6:579–591.

18. Greening C, Ahmed FH, Mohamed AE, Lee BM, Pandey G, Warden AC, Scott C, Oakeshott JG, Taylor MC, Jackson CJ. 2016. Physiology, Biochemistry, and Applications of F420- and Fo-Dependent Redox Reactions. Microbiol Mol Biol Rev 80:451–493.

19. Jirapanjawat T, Ney B, Taylor MC, Warden AC, Afroze S, Russell RJ, Lee BM, Jackson CJ, Oakeshott JG, Pandey G, Greening C. 2016. The Redox Cofactor F420 Protects Mycobacteria from Diverse Antimicrobial Compounds and Mediates a Reductive Detoxification System. Appl Environ Microbiol 82:6810–6818.

20. Purwantini E, Mukhopadhyay B. 2013. Rv0132c of Mycobacterium tuberculosis Encodes a Coenzyme F420-Dependent Hydroxymycolic Acid Dehydrogenase. PLoS One 8:e81985.

21. Wang P, Bashiri G, Gao X, Sawaya MR, Tang Y. 2013. Uncovering the enzymes that catalyze the final steps in oxytetracycline biosynthesis. J Am Chem Soc 135:7138–7141.

22. Peschke U, Schmidt H, Zhang HZ, Piepersberg W. 1995. Molecular characterization of the lincomycin-production gene cluster of Streptomyces lincolnensis 78-11. Mol Microbiol 16:1137–1156.

23. Purwantini E, Mukhopadhyay B. 2009. Conversion of NO2 to NO by reduced coenzyme F420 protects mycobacteria from nitrosative damage. Proc Natl Acad Sci U S A 106:6333– 6338.

24. Rifat D, Li S-Y, Ioerger T, Shah K, Lanoix J-P, Lee J, Bashiri G, Sacchettini J, Nuermberger E. 2020. Mutations in fbiD (Rv2983) as a Novel Determinant of Resistance to Pretomanid and Delamanid in Mycobacterium tuberculosis. Antimicrob Agents Chemother 65:e01948–20.

25. Guerra-Lopez D, Daniels L, Rawat M. 2007. Mycobacterium smegmatis mc2 155 fbiC and MSMEG_2392 are involved in triphenylmethane dye decolorization and coenzyme F420 biosynthesis. Microbiology 153:2724–2732.

26. Darwin KH, Ehrt S, Gutierrez-Ramos J-C, Weich N, Nathan CF. 2003. The Proteasome of Mycobacterium tuberculosis Is Required for Resistance to Nitric Oxide. Science 302:1963– 1966.

27. Rupasinghe P, Ismail N, Mulders W, Warren RM, Joseph L, Ngcamu D, Gwala T, Omar SV, Vereecken J, de Jong BC, Rigouts L, Borroni E, Cirillo DM, Schön T, Miotto P, Köser CU. In vitro exposure to clofazimine can select for delamanid and pretomanid resistance in Mycobacterium tuberculosis. Antimicrob Agents Chemother 69:e01113–25.

28. Ismail N, Dippenaar A, Warren RM, Peters RPH, Omar SV. Emergence of Canonical and Noncanonical Genomic Variants following In Vitro Exposure of Clinical Mycobacterium tuberculosis Strains to Bedaquiline or Clofazimine. Antimicrob Agents Chemother 67:e01368–22.

29. Waller NJE, Cheung C-Y, Cook GM, McNeil MB. 2023. The evolution of antibiotic resistance is associated with collateral drug phenotypes in Mycobacterium tuberculosis. Nat Commun 14:1517.

30. Jirapanjawat T, Ney B, Taylor MC, Warden AC, Afroze S, Russell RJ, Lee BM, Jackson CJ, Oakeshott JG, Pandey G, Greening C. 2016. The Redox Cofactor F420 Protects Mycobacteria from Diverse Antimicrobial Compounds and Mediates a Reductive Detoxification System. Appl Environ Microbiol 82:6810–6818.

31. Malachite Green - an overview | ScienceDirect Topics. https://www.sciencedirect.com/topics/agricultural-and-biological-sciences/malachite-green. Retrieved 23 May 2025.

32. Selengut JD, Haft DH. 2010. Unexpected Abundance of Coenzyme F420-Dependent Enzymes in Mycobacterium tuberculosis and Other Actinobacteria. J Bacteriol 192:5788– 5798.

33. Cellitti SE, Shaffer J, Jones DH, Mukherjee T, Gurumurthy M, Bursulaya B, Boshoff HI, Choi I, Nayyar A, Lee YS, Cherian J, Niyomrattanakit P, Dick T, Manjunatha UH, Barry CE, Spraggon G, Geierstanger BH. 2012. Structure of Ddn, the Deazaflavin-Dependent Nitroreductase from Mycobacterium tuberculosis Involved in Bioreductive Activation of PA-824. Structure 20:101–112.

34. Purwantini E, Daniels L. 1998. Molecular Analysis of the Gene Encoding F420-Dependent Glucose-6-Phosphate Dehydrogenase fromMycobacterium smegmatis. Journal of Bacteriology 180:2212–2219.

35. Aderemi AV, Snee M, Tunnicliffe RB, Golovanova M, Cain KM, Munro AW, Waltho JP, Leys D. 2025. Expression and purification of *Mycobacterium tuberculosis* F420-dependent glucose-6-phosphate dehydrogenase enzyme using *Escherichia coli*. Protein Expression and Purification 228:106650.

36. Hasan MR, Rahman M, Jaques S, Purwantini E, Daniels L. 2010. Glucose 6-Phosphate Accumulation in Mycobacteria: IMPLICATIONS FOR A NOVEL F420-DEPENDENT ANTI-OXIDANT DEFENSE SYSTEM *. Journal of Biological Chemistry 285:19135–19144.

37. Matern WM, Harris HT, Danchik C, McDonald M, Patel G, Srivastava A, Ioerger TR, Bader JS, Karakousis PC. 2023. Functional Whole Genome Screen of Nutrient-Starved Mycobacterium tuberculosis Identifies Genes Involved in Rifampin Tolerance. Microorganisms 11:2269.

38. Gopal P, Tasneen R, Yee M, Lanoix J-P, Sarathy J, Rasic G, Li L, Dartois V, Nuermberger E, Dick T. 2017. In Vivo-Selected Pyrazinoic Acid-Resistant Mycobacterium tuberculosis Strains Harbor Missense Mutations in the Aspartate Decarboxylase PanD and the Unfoldase ClpC1. ACS Infect Dis 3:492–501.

39. Almeida D, Converse PJ, Li S-Y, Upton AM, Fotouhi N, Nuermberger EL. 2021. Comparative Efficacy of the Novel Diarylquinoline TBAJ-876 and Bedaquiline against a Resistant Rv0678 Mutant in a Mouse Model of Tuberculosis. Antimicrob Agents Chemother 65:e0141221.

40. Lamichhane G, Zignol M, Blades NJ, Geiman DE, Dougherty A, Grosset J, Broman KW, Bishai WR. 2003. A postgenomic method for predicting essential genes at subsaturation levels of mutagenesis: Application to Mycobacterium tuberculosis. Proc Natl Acad Sci U S A 100:7213–7218.

41. Rifat D, Chen L, Kreiswirth BN, Nuermberger EL. 2021. Genome-Wide Essentiality Analysis of Mycobacterium abscessus by Saturated Transposon Mutagenesis and Deep Sequencing. mBio 12:e0104921.

42. Long JE, DeJesus M, Ward D, Baker RE, Ioerger T, Sassetti CM. 2015. Identifying Essential Genes in Mycobacterium tuberculosis by Global Phenotypic Profiling, p. 79–95. In Lu, LJ (ed.), Gene Essentiality: Methods and Protocols. Springer, New York, NY.

43. DeJesus MA, Ambadipudi C, Baker R, Sassetti C, Ioerger TR. 2015. TRANSIT - A Software Tool for Himar1 TnSeq Analysis. PLoS Comput Biol 11:e1004401.

44. DeJesus MA, Ioerger TR. 2016. NORMALIZATION OF TRANSPOSON-MUTANT LIBRARY SEQUENCING DATASETS TO IMPROVE IDENTIFICATION OF CONDITIONALLY ESSENTIAL GENES. J Bioinform Comput Biol 14:1642004.

45. Lun S, Guo H, Onajole OK, Pieroni M, Gunosewoyo H, Chen G, Tipparaju SK, Ammerman NC, Kozikowski AP, Bishai WR. 2013. Indoleamides are active against drug-resistant Mycobacterium tuberculosis. Nat Commun 4:2907.

46. Briffotaux J, Huang W, Wang X, Gicquel B. 2017. MmpS5/MmpL5 as an efflux pump in *Mycobacterium* species. Tuberculosis 107:13–19.

47. Radhakrishnan A, Kumar N, Wright CC, Chou T-H, Tringides ML, Bolla JR, Lei H-T, Rajashankar KR, Su C-C, Purdy GE, Yu EW. 2014. Crystal Structure of the Transcriptional Regulator Rv0678 of *Mycobacterium tuberculosis*\*. Journal of Biological Chemistry 289:16526–16540.

48. Ioerger TR, O’Malley T, Liao R, Guinn KM, Hickey MJ, Mohaideen N, Murphy KC, Boshoff HIM, Mizrahi V, Rubin EJ, Sassetti CM, Iii CEB, Sherman DR, Parish T, Sacchettini JC. 2013. Identification of New Drug Targets and Resistance Mechanisms in Mycobacterium tuberculosis. PLOS ONE 8:e75245.

49. Zhang S, Chen J, Cui P, Shi W, Zhang W, Zhang Y. 2015. Identification of novel mutations associated with clofazimine resistance in Mycobacterium tuberculosis. J Antimicrob Chemother 70:2507–2510.

50. Hartkoorn RC, Uplekar S, Cole ST. 2014. Cross-Resistance between Clofazimine and Bedaquiline through Upregulation of MmpL5 in Mycobacterium tuberculosis. Antimicrobial Agents and Chemotherapy 58:2979–2981.

51. Alderman DJ. 1985. Malachite green: a review. Journal of Fish Diseases 8:289–298.

52. Srivastava S, Sinha R, Roy D. 2004. Toxicological effects of malachite green. Aquat Toxicol 66:319–329.

53. Ngcobo BP, Osunsanmi FO, Ikhane AO, Mosa RA, Opoku AR. 2025. Biodegradation and detoxification of malachite green dye by crude manganese peroxidase produced by Pseudomonas aeruginosa MF14446.1 and Enterobacter roggenkampii CP033800.1. Sci Rep 10.1038/s41598-025-33918-3.

54. Wang J, Gao F, Liu Z, Qiao M, Niu X, Zhang K-Q, Huang X. 2012. Pathway and Molecular Mechanisms for Malachite Green Biodegradation in Exiguobacterium sp. MG2. PLoS ONE 7:e51808.

55. Jang M-S, Lee Y-M, Kim C-H, Lee J-H, Kang D-W, Kim S-J, Lee Y-C. 2005. Triphenylmethane Reductase from Citrobacter sp. Strain KCTC 18061P: Purification, Characterization, Gene Cloning, and Overexpression of a Functional Protein in Escherichia coli. Appl Environ Microbiol 71:7955–7960.

56. Azmi W, Sani RK, Banerjee UC. 1998. Biodegradation of triphenylmethane dyes. Enzyme and Microbial Technology 22:185–191.

57. Gu S, Chen J, Dobos KM, Bradbury EM, Belisle JT, Chen X. 2003. Comprehensive proteomic profiling of the membrane constituents of a Mycobacterium tuberculosis strain. Mol Cell Proteomics 2:1284–1296.

58. Mawuenyega KG, Forst CV, Dobos KM, Belisle JT, Chen J, Bradbury EM, Bradbury ARM, Chen X. 2005. Mycobacterium tuberculosis functional network analysis by global subcellular protein profiling. Mol Biol Cell 16:396–404.

59. Ouellet H, Johnston JB, Ortiz de Montellano PR. 2010. The Mycobacterium tuberculosis Cytochrome P450 System. Arch Biochem Biophys 493:82–95.

60. Lun S, Bishai WR. 2007. Characterization of a Novel Cell Wall-anchored Protein with Carboxylesterase Activity Required for Virulence in *Mycobacterium tuberculosis*\*. Journal of Biological Chemistry 282:18348–18356.

61. Naffin-Olivos JL, Daab A, White A, Goldfarb NE, Milne AC, Liu D, Baikovitz J, Dunn BM, Rengarajan J, Petsko GA, Ringe D. 2017. Structure Determination of Mycobacterium tuberculosis Serine Protease Hip1 (Rv2224c). Biochemistry 56:2304–2314.

62. Lun S, Miranda D, Kubler A, Guo H, Maiga MC, Winglee K, Pelly S, Bishai WR. 2014. Synthetic Lethality Reveals Mechanisms of Mycobacterium tuberculosis Resistance to β-Lactams. mBio 5:e01767–14.

63. Xu W, DeJesus MA, Rücker N, Engelhart CA, Wright MG, Healy C, Lin K, Wang R, Park SW, Ioerger TR, Schnappinger D, Ehrt S. 2017. Chemical Genetic Interaction Profiling Reveals Determinants of Intrinsic Antibiotic Resistance in Mycobacterium tuberculosis. Antimicrob Agents Chemother 61:e01334–17.

64. Rengarajan J, Murphy E, Park A, Krone CL, Hett EC, Bloom BR, Glimcher LH, Rubin EJ. 2008. Mycobacterium tuberculosis Rv2224c modulates innate immune responses. Proceedings of the National Academy of Sciences 105:264–269.

65. Almeida DV, Converse PJ, Nuermberger EL. Mutations in Rv0678 Reduce Susceptibility of Mycobacterium tuberculosis to the DprE1 Inhibitor TBA-7371. Antimicrob Agents Chemother 67:e00052–23.

66. Poulton NC, Azadian ZA, DeJesus MA, Rock JM. Mutations in rv0678 Confer Low-Level Resistance to Benzothiazinone DprE1 Inhibitors in Mycobacterium tuberculosis. Antimicrob Agents Chemother 66:e00904–22.

67. Hariguchi N, Chen X, Hayashi Y, Kawano Y, Fujiwara M, Matsuba M, Shimizu H, Ohba Y, Nakamura I, Kitamoto R, Shinohara T, Uematsu Y, Ishikawa S, Itotani M, Haraguchi Y, Takemura I, Matsumoto M. 2020. OPC-167832, a Novel Carbostyril Derivative with Potent Antituberculosis Activity as a DprE1 Inhibitor. Antimicrob Agents Chemother 64:e02020–19.

68. Boshoff HIM, Myers TG, Copp BR, McNeil MR, Wilson MA, Barry CE. 2004. The Transcriptional Responses of *Mycobacterium tuberculosis* to Inhibitors of Metabolism. Journal of Biological Chemistry 279:40174–40184.

69. Imlay JA. 2008. Cellular defenses against superoxide and hydrogen peroxide. Annu Rev Biochem 77:755–776.

70. Cumming BM, Lamprecht DA, Wells RM, Saini V, Mazorodze JH, Steyn AJC. 2014. The Physiology and Genetics of Oxidative Stress in Mycobacteria. Microbiology Spectrum 10.1128/microbiolspec.mgm2-0019–2013.

71. MacMicking J, Xie Q, Nathan C. 1997. NITRIC OXIDE AND MACROPHAGE FUNCTION. Annu Rev Immunol 15:323–350.

72. Jamaati H, Mortaz E, Pajouhi Z, Folkerts G, Movassaghi M, Moloudizargari M, Adcock IM, Garssen J. 2017. Nitric Oxide in the Pathogenesis and Treatment of Tuberculosis. Front Microbiol 8:2008.

73. Prosser G, Brandenburg J, Reiling N, Barry CE, Wilkinson RJ, Wilkinson KA. 2017. The bacillary and macrophage response to hypoxia in tuberculosis and the consequences for T cell antigen recognition. Microbes Infect 19:177–192.

74. Liu A. 2024. Catalase-peroxidase (KatG): a potential frontier in tuberculosis drug development. Critical Reviews in Biochemistry and Molecular Biology 59:434–446.

75. Nambi S, Long JE, Mishra BB, Baker R, Murphy KC, Olive AJ, Nguyen HP, Shaffer SA, Sassetti CM. 2015. The Oxidative Stress Network of *Mycobacterium tuberculosis* Reveals Coordination between Radical Detoxification Systems. Cell Host & Microbe 17:829–837.

76. Rachman H, Strong M, Ulrichs T, Grode L, Schuchhardt J, Mollenkopf H, Kosmiadi GA, Eisenberg D, Kaufmann SHE. 2006. Unique Transcriptome Signature of Mycobacterium tuberculosis in Pulmonary Tuberculosis. Infect Immun 74:1233–1242.

77. Sherman DR, Voskuil M, Schnappinger D, Liao R, Harrell MI, Schoolnik GK. 2001. Regulation of the Mycobacterium tuberculosis hypoxic response gene encoding α-crystallin. Proc Natl Acad Sci U S A 98:7534–7539.

78. Awuh JA, Flo TH. 2016. Molecular basis of mycobacterial survival in macrophages. Cell Mol Life Sci 74:1625–1648.

79. Bellerose MM, Proulx MK, Smith CM, Baker RE, Ioerger TR, Sassetti CM. 2020. Distinct Bacterial Pathways Influence the Efficacy of Antibiotics against Mycobacterium tuberculosis. mSystems 5:e00396–20.

